# Apical extracellular matrix regulates fold morphogenesis in the *Drosophila* wing disc

**DOI:** 10.1101/2025.09.06.674631

**Authors:** Jana F. Fuhrmann, Vincenzo Maria Schimmenti, Greta Cwikla, Sangwon Lee, Michaela Yuan, Michaela Wilsch-Bräuninger, Frank Jülicher, Marko Popović, Natalie A. Dye

## Abstract

Tissue folding is a fundamental process occurring often in animal organ development. Here, we study the progression of fold shape and the underlying mechanics in the development of the *Drosophila* wing disc. We present a 3D segmentation of the apical surface of the wing disc proper from larval stages, when folds grow, to early pupariation, when the tissue unfolds and remodels into a bilayer. We establish morphological metrics to quantify and resolve fold shape in this dataset, introducing a definition of fold depth and width that can be used to characterize folds on a curved surface. Furthermore, we identify fibrous extracellular matrix on the apical side (aECM) that physically connects the two opposing sides of the folds. By modeling a tissue fold with a lateral vertex model endowed by an adhesive layer representing the aECM, we predict that unfolding in the wing disc is preceded by the removal of aECM. Using genetic perturbations, we confirm that aECM adhesion affects fold stability and mechanics: loss of aECM leads to abnormal fold shape and unfolding dynamics, whereas failure to remove aECM at pupariation inhibits unfolding. Finally, we show that these aECM perturbations in larval stages cause morphological phenotypes in the adult wing, demonstrating that the fold morphology of the wing disc helps to define adult wing shape. In total, our work establishes a key mechanical role for aECM in wing disc growth and morphogenesis and advances our general understanding of how epithelial tissue folds can be mechanically stabilized during development.

## Introduction

Understanding how an animal tissue can grow into a particular 3D shape and reshape itself dynamically during its development remains a major goal of developmental biology and biophysics. In particular, tissue folds are a common feature of 3D tissue architecture. Folds can become part of the final organ structure, such as those of the human brain, or serve as a transient morphogenetic process, such as ventral furrow formation in the gastrulating *Drosophila* embryo [1].

To date, multiple mechanisms have been described that lead to folding in different tissues (as reviewed in [2, 3]). For example, apical constriction generated by actin-myosin in epithelial cells changes the cell aspect ratio to induce bending in various contexts. Folding can also occur through buckling induced by differential growth within the epithelium or between neighboring tissues [2, 3].

Little is known about how folds propagate, mature, and stabilize during growth and homeostasis. Here, we study the folds of the *Drosophila* wing disc, which are a stereotypic set of transient folds that grow over 2-3 days, before opening up over hours during metamorphosis [4, 5]. This developmental trajectory begs the question of what stabilizes these folds during the growth phase, how they later become destabilized, and what is the morphogenetic consequence of this sequence for wing development. The wing disc folds thus provide an excellent system to address the open general topic of how folds propagate in depth, width, and length in a growing epithelium and how the mechanical state of the folds mediates stability.

The *Drosophila* larval wing disc is composed of two opposing epithelial cell layers that face apical inwards. The pseudostratified, columnar wing disc proper layer gives rise to the adult wing, hinge, and notum, while the other layer, the peripodial epithelium (PE), is squamous and is removed at metamorphosis. The wing disc is connected only at its proximal-most region to the larval epithelium by the stalk cells. Thus, the majority of wing disc tissue does not have direct physical interaction with other tissues, and its mechanics can be assumed to be self-contained.

The wing disc epithelium is surrounded by extracellular matrix on the apical and basal side [6, 7, 8, 9, 10]. The basal ECM contains conserved ECM proteins, including Collagen-IV and laminin, and has many known roles for maintaining wing disc architecture [7, 11, 12, 13, 14]. The composition of apical ECM (aECM) in the wing disc is less clear but contains at least the Dumpy protein, a giant protein with hundreds of epidermal growth factor modules and a Zona Pellucida (ZP) domain [15, 9, 10]. Generally in *Drosophila*, the composition and structure of aECM can vary across tissues, but ZP-domain proteins are a common component. They also form a link between the apical membrane and the aECM ([16]). Importantly, the function of aECM in disc proper morphogenesis is unclear.

The wing disc grows from a flat disc into a complex, folded 3D structure over approximately three days [4]. The most prominent and well-studied folds are the three folds across the dorsal side of the epithelium in the Proximal-Distal (PD) axis that morphologically separate the presumptive adult body parts. The dome-like pouch structure gives rise to the distal proportion of the adult wing blade and is separated by a fold (hinge-pouch, HP fold) from more proximal regions that contribute to the proximal wing blade. The wing hinge forms from the intermediate region and is divided by a central fold (hinge-hinge, HH fold). The third and most proximal fold separates the prospective notum from the wing hinge (hinge-notum, HN fold). Notably, previous works have identified different mechanisms for fold initiation [12, 17], whereas the question of how fold morphology evolves and what is the role of the folds during wing disc development remains open.

The vast majority of growth and patterning of the wing happens during larval stages, and therefore perturbations of these processes at this stage cause adult phenotypes. Nevertheless, there is additionally considerable remodeling of the tissue during pupal stages that also affects adult morphology [18, 19, 9]. At the onset of pupariation, the wing disc unfolds and everts, transforming into the flat bilayered wing and the future hinge and notum. During this process, the formation of the bilayer is achieved through an anisotropic expansion by spatially patterned cell rearrangements and cell area changes in the pouch [5]. Simultaneously the PE gets removed, mediated by actin-myosin contraction [20], and the wing disc is placed outside of the body [21]. Later, the presumptive wing blade region undergoes columnar-to-cuboidal shape changes and elongates [8]. These processes are mediated by a reorganization of actin-myosin and ECM removal [8, 22]. Immediately afterwards, new connections are established between the opposing basal sides of the epithelial bilayer that are required for a flat adult wing shape [23].

Here, we study how fold morphology in the developing wing disc evolves in time, how the aECM contributes to these dynamics, and how early folding dynamics affect adult wing morphology. We first provide a 3D quantification of the spatial and temporal geometry of the folds during growth and eversion, going beyond the standard of analyzing folds in single cross-sections. To do so, we created triangulated meshes of the wing disc apical surface throughout development and generated an analysis method to quantify fold morphology. Using ultrastructure analysis and genetic reporter lines, we identify aECM as the molecular scaffold that connects the apical sides of the wing disc folds. We then use a physical model of a wing disc fold with aECM represented by adhesive interaction between apposing cell layers in the fold. This model reveals different modes of fold opening dynamics when aECM is removed or remains present. Comparison of model results to experimental data suggests that aECM is biochemically removed at the onset of eversion. We test the role of aECM in fold morphogenesis using genetic perturbations, either removing aECM prematurely or preventing aECM removal at the onset of eversion. Our results show that aECM is critical for fold shape and stability during growth and that removal of aECM is critical for wing disc unfolding and to obtain a flat adult fly wing. Overall, our work establishes aECM as an important contribution to wing disc tissue mechanics that regulates fold morphogenesis and that folding dynamics are important for correct adult wing morphology.

## Results

### 3D imaging and quantification reveal fold dynamics throughout development

We first sought to characterize the morphological changes of the wing disc proper during larval stages and wing disc eversion. To this end, we explanted live wing discs at different developmental stages and used multi-angle light sheet microscopy to attain a full 3D description of the tissue. Our data includes three growth time points, each separated by one day, followed by the last larval wing disc shape (wandering 3rd instar, wL3) which marks the end of larval development and onset of wing eversion. We also include eversion time points, starting from 0 hours after puparium formation (hAPF) to 6 hAPF, when the folds are open and the tissue resembles a flat bilayer.

To quantify the morphological complexity of the wing disc, we segmented the E-Cadherin::GFP (Ecad::GFP) signal on the apical surface of the wing disc proper and generated surface meshes from the basal-facing edge of the segmentation. The sub-apical mesh (below adherens junctions) allows for the separation of the apposing tissue surfaces in tightly folded regions (Figure 1A and Figure S1A,B). The resulting 3D apical surface description shows the shape changes in larval stages, during which the wing disc grows and transforms from a flat monolayer geometry to a larger highly folded structure over the time course of 2-3 days, from 3 days after egg laying (dAEL) to wL3 stage (Figure 1B-C’ and Figure S1C,D). We color the meshes by mean curvature, where we choose the sign of curvature to be positive in folds and negative on ridges. These data confirm the previously reported sequential formation of the folds during growth, starting with the HH fold and followed by the HN and HP folds [12]. Interestingly, our data also reveal the emergence of folds on the lateral and ventral sides of the wing pouch, which are often overlooked (Figure 1B-C’ and Figure S1C,D). We also show the process of unfolding, bilayer formation, and tissue flattening during wing eversion from wL3 to 6 hAPF which is completed over a fast time course in the range of 10-12 h, estimating 4-6 h for wL3 (Figure 1C,C’ and Figure S1C,D) [5]

**Figure 1:**
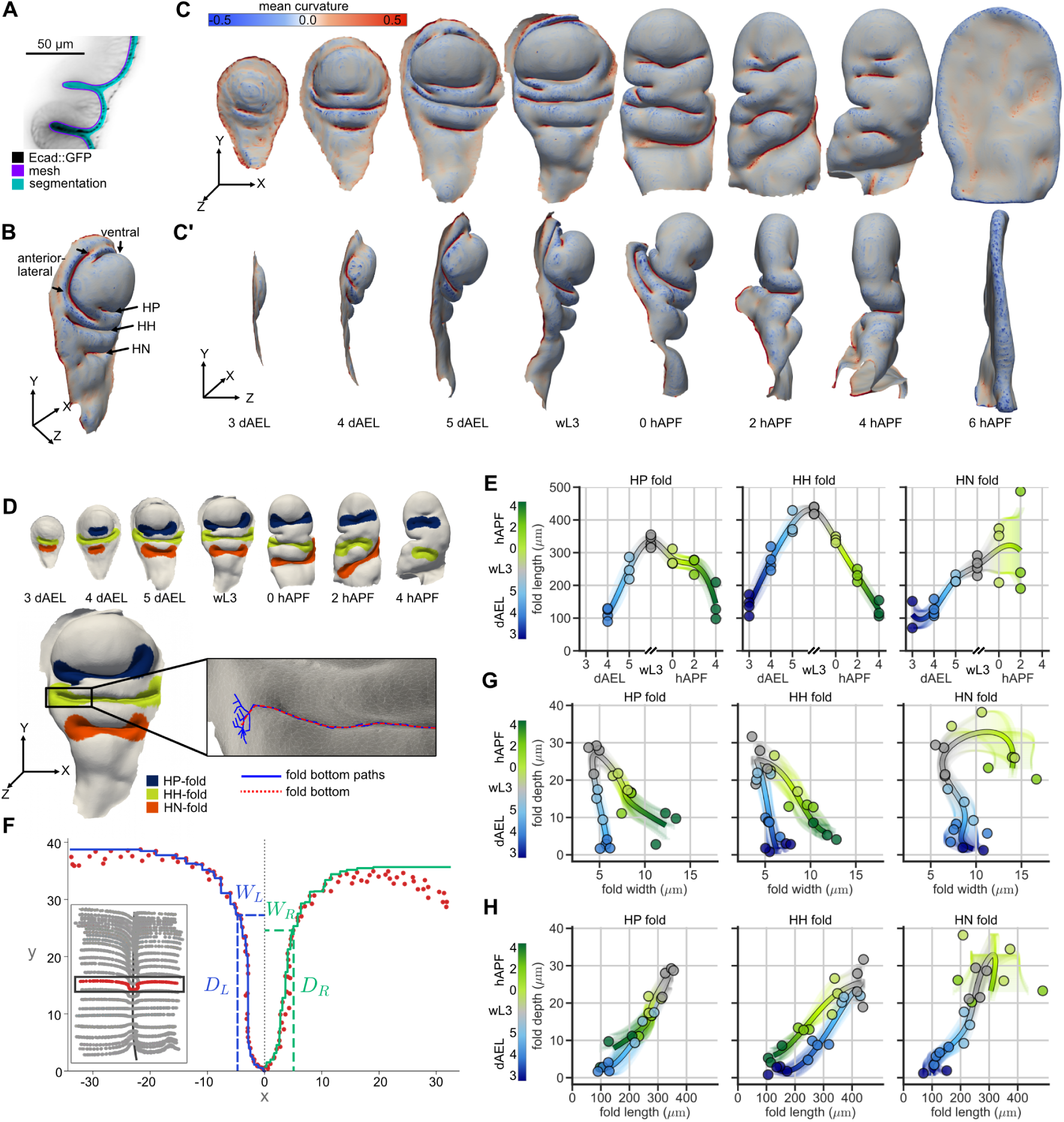
**A**, Example cross-section of a 5 dAEL wing disc showing Ecad::GFP, the apical segmentation result, and the position of the mesh. **B**, Mesh surface of the wing disc from A, with the different folds labeled (HP = hinge-pouch, HH = hinge-hinge, HN = hinge-notum) and mean curvature (*µm^−^*^1^) represented by the color of the mesh faces. **C-C’**, Wing disc proper apical surface meshes with mean curvature (*µm^−^*^1^) represented as in **B**. **C**, Head-on view: Larval stages (3 dAEL - wL3), with anterior left, and ventral pointed up; after bilayer formation (0-6 hAPF), only the dorsal side is visible in this orientation. **C’**, Anterior-lateral view. Wing discs 0 - 6 hAPF are cropped on the notum. **D**, Top: Segmented fold region at different time points. The color corresponds to the fold identity. Below: Identification of the fold bottom curve for the HH fold of a wL3 stage wing disc. The fold bottom curve (red) is identified from candidate fold bottom paths (blue). **E, G, H**, Plots of each fold, with color indicating the developmental time. Data points indicate measured quantities of individual replicates, and the line shows the mean cubic spline interpolation over all data points. Error bars show the standard deviation of the mean spline. **E**, Fold length over developmental time. **F**, Plot of an individual fold cross-section, where red points correspond to the mesh vertices and the solid lines show the left and right envelope. Dashed lines identify width and depth for each envelope. Inset: all cross-sections of a fold bottom curve (grey), with the individual cross-section highlighted in red. **G**, Plot of fold depth and width, colored by developmental time. **H**, Plot of fold depth and length, colored by developmental time.

To characterize the dynamics of folding and unfolding, we established the methodology to identify the folds. In particular, the fold regions are characterized by a typical high positive mean curvature, and folds are separated by ‘hills’ with a negative mean curvature. This allows us to define the fold regions based on curvature thresholding (Methods). While this method identifies all folds in the tissue, we focus our analysis on the shape of the HP, HH, and HN folds from 3 dAEL to 4 hAPF. To characterize fold morphology, we use familiar quantities such as length, depth, and width. Fold length specifies the length of the long axis, which for the HP, HH, and HN folds corresponds roughly to the anterior-posterior axis of the wing disc. Width and depth are specified in a cross-section orthogonal to the long axis. Since folds are narrow and long but change their direction and cross-section, we first needed to define a reference frame inside the fold. To this end, we identify a fold bottom curve, which runs along the highest mean curvature of the segmented fold region. As the curvature decreases near the ends of the folds, this bottom curve becomes less well defined. Therefore, we merge multiple candidate curves starting from different possible endpoints using a dynamic time warping algorithm, see Methods and Figure 1D.

In this way, we measure the length of the folds over time. At 3 dAEL, the HP fold is not identifiable with this method (Figure 1D), supporting our observation that it forms after the HH and HN folds. Moreover, at 4 hAPF, the wing disc is much larger, and the HN fold is often outside the field of view of our images. Thus, we restrict our analysis of the HN fold to up to 2 hAPF wing discs. We find that fold length increases during wing disc growth (Figure 1E). In the HP and HH folds, a maximum fold length is reached at wL3, after which the folds shorten as they unfold. In contrast, the HN fold does not undergo fold shortening during eversion (Figure 1E).

To further address the dynamics of fold shape changes, we devised a method to quantify fold width and depth. To this end, we divide the fold into cross-sections along the fold bottom, as shown in Figure 1F and described in Methods. As the bottom curve orientation changes along the fold, we define the local reference frame in each cross-section, see Methods. To take into account asymmetries between two sides of the fold, in each cross-section we first define depth and width on each side of the fold, left and right relative to the fold bottom along the *x*-axis in the local reference frame. For this, we construct an upper envelope of the cross-section points on each side. An example of the envelope is shown in Figure 1F, see Methods for details. The cross-section widths on the left and right sides *W_L,R_* are defined as the average x-coordinate of the envelope, weighted by the local slope, see Methods. The cross-section depths *D_L,R_* are then the *y* coordinates of the envelopes at the cross-section widths *W_L,R_*, as illustrated in Figure 1F and Figure S2A. Finally, the fold depth and width are obtained by an averaging procedure over the two sides and over all cross-sections along the fold, see Methods and Figure S2A-C.

We measure fold depth and width at different developmental stages in the wing disc (Figure 1G). We find that the HP and HH folds have similar dynamics. Fold depth increases up to wL3, with only a slight narrowing as the folds deepen. However, during eversion, from wL3 to 4 hAPF, the folds follow a different trajectory and get simultaneously shallower and wider. This hysteretic behavior suggests that the mechanics of fold opening and deepening are different. The HN fold shows similar dynamics during deepening. However, during unfolding, the fold widens but does not decrease as much in depth, indicating that the HN fold does not fully unfold by 2 hAPF. Lastly, we look at the relationship between fold depth and length. We find that the fold deepens as it gets longer and becomes shallower as it gets shorter (Figure 1H). Interestingly, the HP fold follows the same path in the depth-length plane during during deepening and opening, although it is much wider during the opening (compare Figure 1G and H). A similar trend can be seen in the HH fold, albeit it appears to be slightly shorter at a given depth during fold opening than it is during fold deepening (Figure 1H).

### aECM connects apposing sides of folds during growth and detaches at eversion

To understand how fold morphology is structurally and mechanically controlled, we performed ultrastructure analysis using transmission electron microscopy (TEM). We observed that the extracellular space on the apical side is filled with highly structured aECM (Figure 2A and Figure S3A). In the lateral cross-section along the long axis of the wing disc, we observe aECM covering the epithelium from the top to the bottom of the folds. Fold cells connect to a shared layer of aECM from both sides via electron-dense tips of apical microvilli (Figure 2A,B and Figure S3B). In contrast, the basal ECM does not appear to be in direct contact with both sides of neighboring cell layers (Figure S3C). Inside the folds and in regions of close contact between cells, the aECM appears fibrous or sheet-like, whereas further away from the apical surface the aECM is more homogeneous (Figure 2B.1). At the contact side with apical protrusions, the aECM is denser, and the space between these protrusions is largely free of ECM (Figure 2B.2).

**Figure 2:**
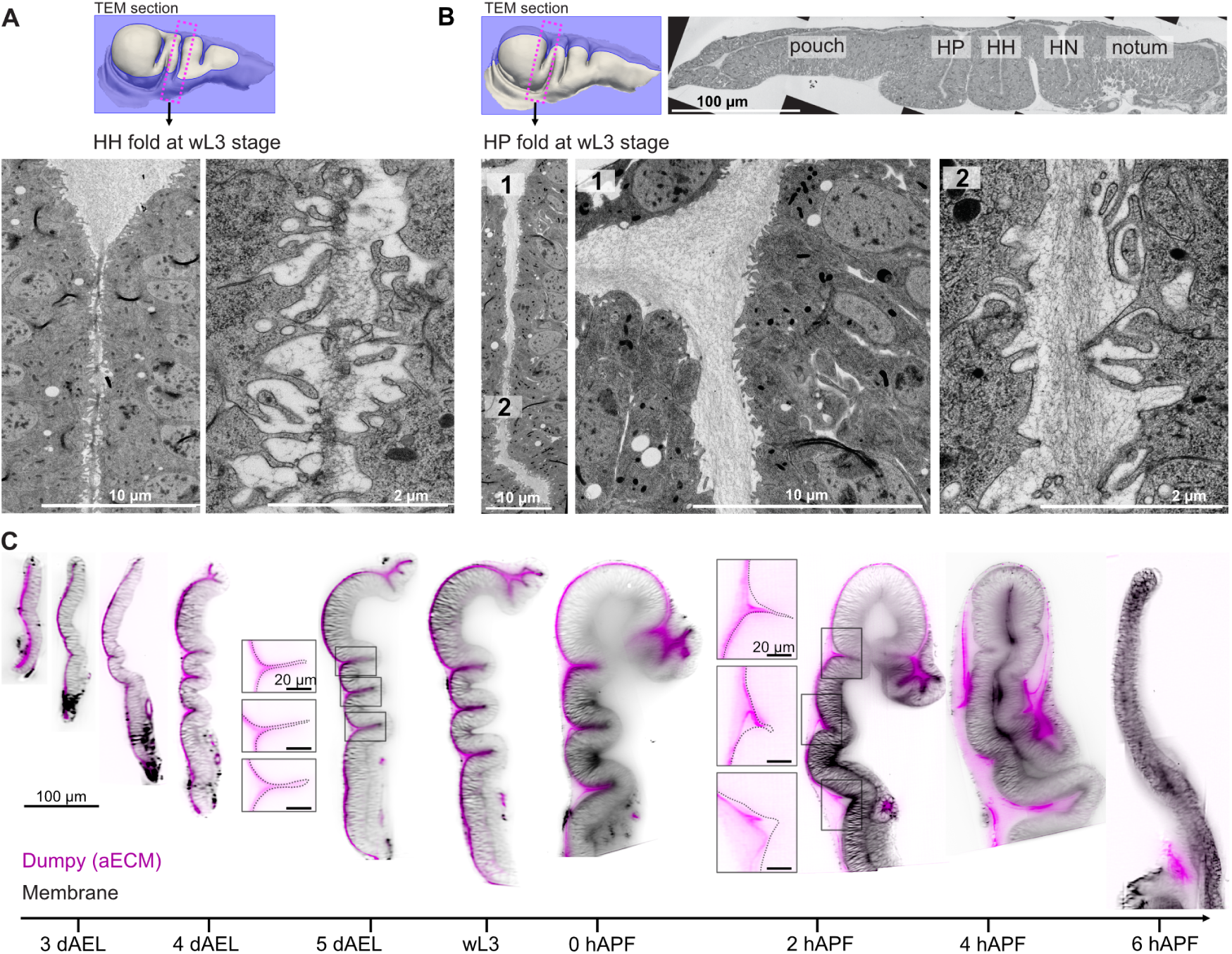
**A,B**, TEM of a wL3 wing disc. Top: Indication of the section orientation: wing disc surface in grey; TEM section in blue. The dashed box (magenta) highlights the region for which the TEM data is shown below. **A**, TEM of the HH fold. Left: overview of the extracellular space and cells of the hinge region. Right: zoom-in of the same region. **B**, TEM of a cross-section along the long axis. Top: overview of the cross-section with the tissue regions labeled. Bottom left: overview of the HP fold, with the position of the higher magnification images shown on the right indicated by numbers. Bottom middle (1): upper HP fold region, showing the apical extracellular space and cells of the PE (top), pouch (left), hinge (right). Bottom right (2): higher magnification of the same fold closer to the fold bottom. **C**, Long-axis cross-sections of light-sheet images of Dumpy::YFP wing discs. Developmental time points are indicated below. Magenta: aECM (Dumpy::YFP); Black: cell membranes (fm464). All wing disc images are to scale, with the scale bar on the left. The insets show higher magnification images of Dumpy::YFP for 120 hAEL and 2 hAPF (scale bars = 20 *µm*). The apical outline of the tissue is indicated by a dashed line. The positions of the insets are indicated by black boxes.

To test the dynamics and abundance of aECM throughout development, we investigated the distribution of Dumpy. Using a Dumpy::YFP fusion, we confirm that aECM covers the entire apical space between peripodial epithelium and disc proper in larval stages and observe that aECM is present throughout fold deepening, suggesting that fold deepening occurs simultaneously with the deposition or restructuring of aECM. The signal intensity is strongest near the disc proper, validating our EM observations that the ECM is denser near the apical outline of the disc proper (see inset Figure 2C, 5 dAEL). During eversion, a gap appears between the Dumpy label and the apical cell surface, indicating that the aECM detaches from the tissue surface. The detachment can be first observed near the opening of the folds from 0 hAPF and continues throughout the disc until all aECM is removed at 6 hAPF (Figure 2C).

### Lateral vertex model with adhesion predicts distinct modes of unfolding depending on aECM degradation

We hypothesized that the aECM has an adhesive function that participates in fold mechanics and that its removal plays a role in successful wing eversion. To study the adhesive role of aECM, we extended the standard lateral vertex model by introducing adhesion between the apical sides of the cells on the apposing sides of the fold, representing the interactions mediated by aECM. We derived an analytical expression for the adhesion energy on the cell scale from a microscopic interaction potential that depends only on the coordinates of cell vertices. This allows for efficient evaluation of the adhesion forces without numerical discretization of the cell surface (Figure 3A).

**Figure 3:**
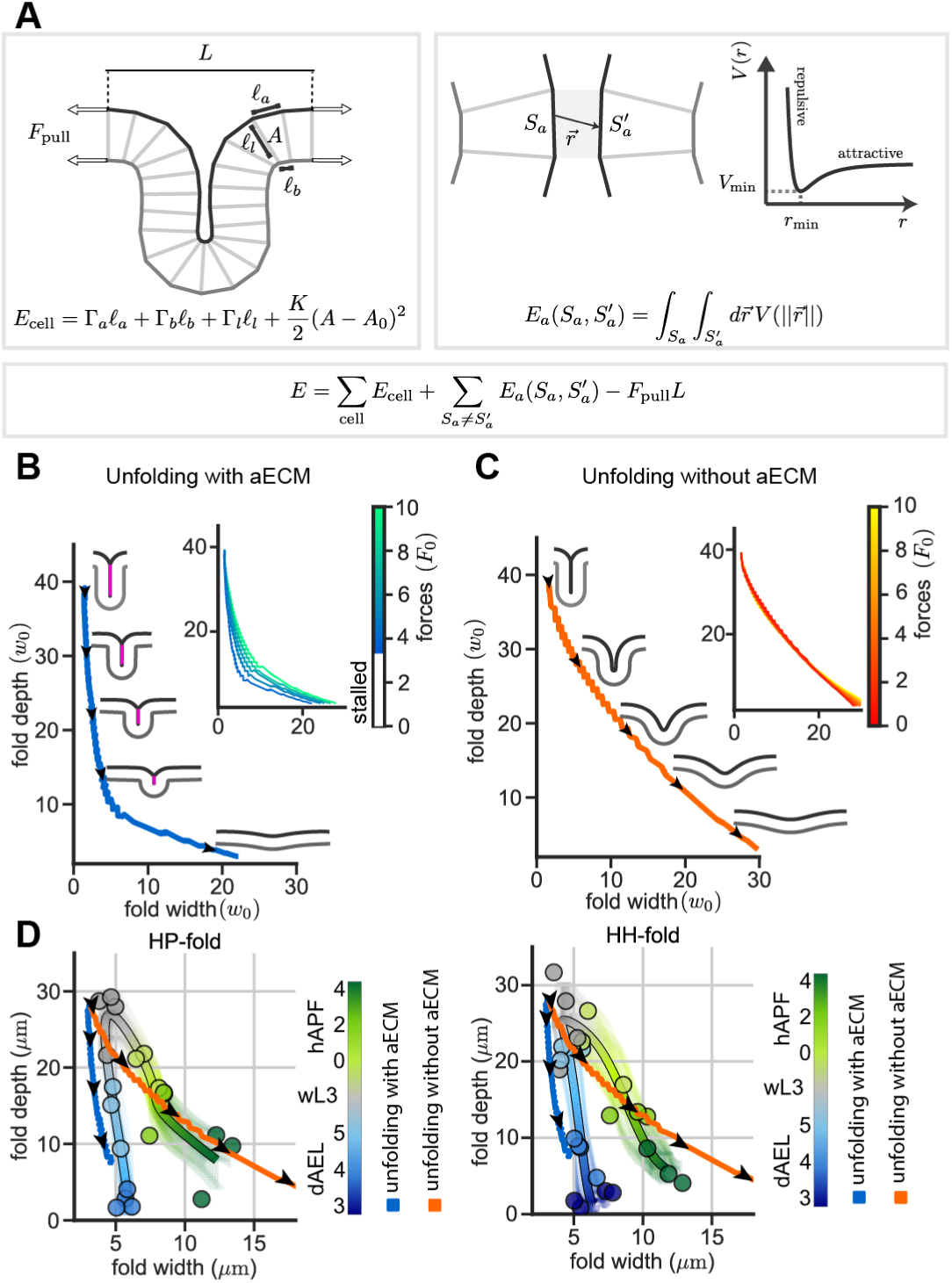
Lateral vertex model of wing unfolding considering aECM adhesion between folds. **A**, Schematic representation of the lateral vertex model and the equal and opposite pulling forces applied at the boundaries. Outermost lateral bonds on each side of the tissue are constrained to be in the vertical position. The lateral vertex model energy function *E*cell is extended with the adhesion energy between pairs of apical cell surfaces *S_a_* and *S^’^_a_*, where the sum is taken over all pairs of apical surfaces whose distance is less than 1.0 (order of width of initial cell, see SI). Adhesion energy is obtained from a generalized Lennard-Jones potential *V* (*||r⃗||*), with the minimum located at *r*min = 0.2*w*0 (see SI) **B,C**, Simulation of fold opening trajectories in the width-depth plane for force *F*pull = 4*F*0. The black arrows show the direction of the shape trajectory, and the inset shows trajectories for different values of pulling forces. Stalled trajectories are not shown, as they are hardly visible. **B**, Simulation with adhesion energy present (depicted by a magenta line). **C**, Simulation of fold opening when the adhesion energy is removed, representing removal of aECM at the onset of eversion, at the same time as the pulling forces are applied. Units of length and force are *w*0 and *F_0_ = 2Kw^3^_0_*, see SI. **D**, Comparison of HP and HH experimental and model fold shape trajectories. Blue and orange curves correspond to model unfolding trajectories with and without aECM, respectively. Units of length in the model are chosen to match the initial fold depth in the model with the depth of wL3 wing disc folds. Model fold trajectories here emulate the slightly sub-apical segmentation of the fold surface in experimental data, as described in SI.

We prepared a folded state of the model tissue to represent the wL3 stage of wing development, as shown in Figure 3A and Figure S4. In this state, the fold shape is kept in equilibrium by apical adhesion and a compressive force *F_eq_* applied to its boundaries. We first wondered whether the wing folds are being opened by a mechanical force originating from surrounding tissues, leading to detachment of the aECM in the process. These forces could arise from active stresses generated in the neighboring tissue regions of the wing disc [5] or in the peripodial epithelium [20]. We simulated this scenario starting from the equilibrium state and applying a boundary pulling force, see Figure 3A. We find that for force magnitudes below the critical value *F ≈* 3*F*_0_ unfolding does not occur, where *F_0_ = 2Kw^3^_0_* is the unit of force in the model, and *w*_0_ is the unit of length, see SI. Beyond the critical value, the tissue unfolds by ‘unzipping’, where fold depth decreases with only a tiny variation in its width, see Figure 3B. This behavior differs from the experimentally observed shape trajectory in the HP and HH folds, where width and depth change simultaneously (Figure 1G). This result suggests that a mechanical pulling force is not sufficient to account for the observed unfolding and that the aECM might need to be removed biochemically.

We therefore implemented a degradation of aECM in our model by removal of the apical adhesion at the same time as the pulling force from the boundary is initiated. We find that the fold widens as it becomes shallower, see Figure 3C, consistent with the observed cross-sectional shapes in the HP and HH folds. Interestingly, for a range of higher pulling force magnitudes, we find that the shape trajectory does not strongly depend on the force magnitude (see inset in Figure 3C). Furthermore, we find that the shape trajectory slope is not sensitive to the initial fold depth as shown in Figure S5.

We directly contrast the two modes of fold opening in the model, with and without aECM removal, with shape trajectories of the HP and HH folds. To do so, we rescale model units to *µm* by matching the initial model fold depth to the wL3 experimental data and taking into account the fact that the segmentation is slightly sub-apical in the experimental data (see SI). We overlay the model and experimental fold shape trajectories in Figure 3D. This comparison highlights how the fold opening trajectory in the wing disc is better recapitulated by the opening trajectory of the model without aECM, in contrast to the model with aECM. In summary, our model of fold unfolding during wing disc eversion points to degradation of aECM as a critical step in the unfolding process and predicts that folds would not be opened without removal of the aECM unless the pulling forces are sufficiently high.

### Protease overexpression removes aECM prematurely and alters fold morphology

The protease Notopleural (Np) is known to target aECM, specifically ZP-domain proteins, and has been shown to be expressed in the wing disc in late larval stages [24, 25]. We generated an inducible Np to manipulate its expression in time and space (Np*^OX^*) and thereby reduce the amount of structured aECM. We combine UAS-Np with a wing disc-specific driver (nub-Gal4) that is strongly expressed throughout wing disc folding and morphogenesis in the distal regions (pouch and HP fold, Figure S6A).

Imaging Np*^OX^* in combination with a Dumpy reporter, we find that the unlike the sheet-like appearance near the apical surface in wild type (WT), the Dumpy signal in Np*^OX^* is homogeneous within the entire apical extracellular space (compare Figure 4A WT and Np*^OX^*). This result suggests that overexpression of Np is sufficient to disrupt the aECM, not only in the region of the nub-Gal4 expression but also in the adjacent regions. To validate that Np*^OX^* perturbs ECM structure, we performed EM imaging on wL3 stage wing discs. We find that with Np*^OX^* the apical protrusions are irregular and the previously observed dense connections between ECM and apical protrusions are absent (Figure 3B and Figure S7A,B). Overall, the aECM is still present but less organized, confirming our live imaging results.

**Figure 4:**
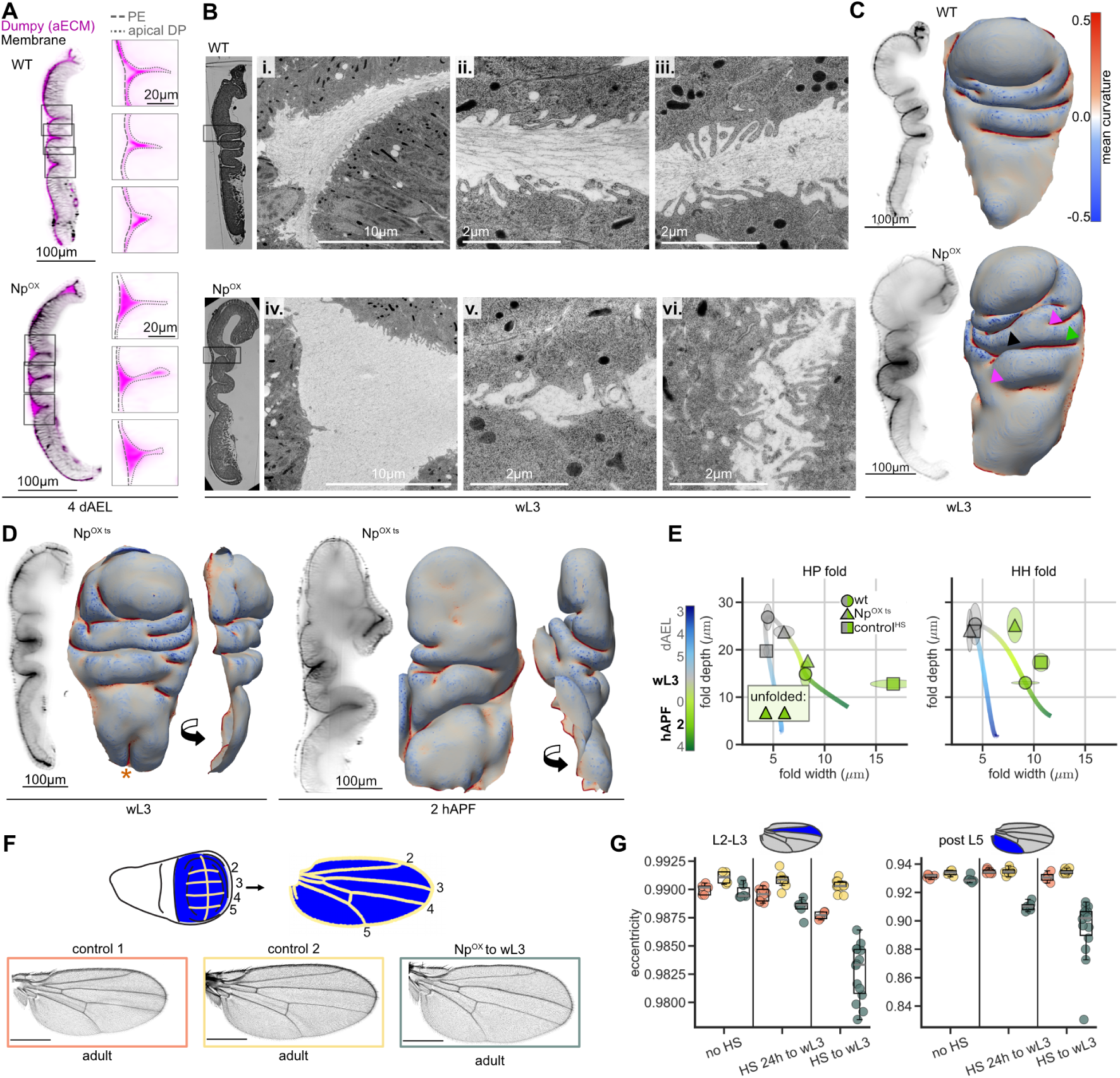
**A**, 4 dAEL wing disc of WT (top) and Np*^OX^* (bottom) in a long-axis cross-section; magenta: aECM (Dumpy::YFP), black: cell membranes (fm464). The insets show higher magnification images of Dumpy::YFP. The apical outline of the cells is indicated by a dashed line (PE = long dashes, DP = short dashes). The positions of the insets are indicated by black boxes. **B**, TEM images of WT (top) and Np*^OX^* (bottom) in a long-axis cross-section. Insets show different positions of the HP-fold. **C,D** Long-axis cross-sections of wing discs labeled with Ecad::GFP and apical surface meshes (color = mean curvature, *µm^−^*^1^) showing head-on and anterior-lateral views. **C**, WT and Np*^OX^* at the wL3 stage. **D**, Np*^OXts^* at wL3 and 2 hAPF stages. **E**, Fold depth over width of HP and HH folds for Np*^OXts^*, WT, and a temperature control at WL3 and 2 hAPF stages, plotted with the WT trajectory. Symbols: mean. Ellipse: the extension along each axis corresponds to the 95% confidence interval of the mean. **F,G**, Analysis of adult wing shape after different temperature shifts. No HS: raised at the permissive temperature for Gal80ts (Gal4 expression should be off) (18°C); 24h to wL3: raised at 18°C, transiently shifted to 30°C (to inhibit Gal80 and allow Gal4 expression) for 24hrs prior to the wL3 stage; to wL3: shifted from egg laying to 30°C and grown there until wL3, whereupon they were shifted and grown thereafter at 18°C. **F**, Cartoon depicting vein regions (yellow) in the wing disc and in the adult fly wing (top) and representative adult wing images of flies grown at 30C to wL3 (bottom). Genotypes are: control1 = nubG4*ts*, control2 = nubG4*ts* x OreR, and Np*^OXts^*. Scale bar = 100 *µm*). **G**, Plots show eccentricity of intervein regions L2-L3 and post L5. The analyzed regions are indicated in blue in the schematic wings above. The genotype is labeled by color as indicated in the colored box around the wings in F.

We then proceed to address the effect of aECM degradation on tissue morphology. We observe a dramatic effect on the shape of the wing disc proper, finding that the anterior lateral fold (Figure 4C, black arrow) extends between the HH and HP folds further than in WT and eventually merges with the HP fold, thereby creating a bifurcated fold (4/4 wing discs, see also Figure S8A,D). A similar effect can be observed at the posterior side, where the posterior lateral fold is longer than in the WT (green arrow). Furthermore, additional bifurcations of folds emerge especially at the posterior side of the HP fold and on the anterior side of the HH fold (magenta arrows, compare also Figure S8C,D). We also see defects in the ventral folds that appear to affect the PE in the surrounding region, consistent with recent work showing that Dumpy knockdown affects cell morphology in the peripodial epithelium [10] (Figure 4C and Figure S8A,B). We quantify the shape of the HH and HN folds and find that the HH fold acquires a similar shape as in the WT, although it is delayed at 5 dAEL, in contrast, the HN fold is shorter, shallower, and wider than in the WT at the wL3 stage (Figure S8D-D”). We see a similar but weaker phenotype with a different Gal4 driver (c765-Np*^OX^*), see Figure S6B and S8E,E’.

We conclude that overexpression of Np throughout the larval development has a dramatic effect on fold shape. It does not, however, inhibit fold formation, nor does it lead to premature fold opening before eversion. Using an overexpression throughout larval stages does not enable us to distinguish the role of aECM during fold growth from its role in fold opening. To specifically test the role of aECM removal for unfolding, we performed a temperature-controlled Np*^OX^* starting 24hrs before wL3 stage, denoted Np*^OXts^*. This setup achieves aECM removal well in advance of eversion, as confirmed by TEM imaging, while reducing any potential influence on fold growth (Figure S7C).

In Np*^OXts^*, we find additional folds, as well as bifurcated folds, similar in structure to folds in the Np*^OX^* experiments described above (Figure 4D and Figure S9A,B). However, the fold pattern is more variable than in Np*^OX^*, and at the proximal tip of the notum we find an additional longitudinal fold (orange asterisk in Figure 4D and Figure S9 compare A and B). We next look at the wing surface shape during eversion at 2 hAPF. We find that the HP fold appears fully unfolded in 2 out of 3 wing discs, while the HH and HN folds are still present (Figure 4D and Figure S9, compare D,D’ and E,E’).

To quantify the unfolding in Np*^OXts^*, we measured the depth and width of the HP and HH folds at wL3 and 2 hAPF. In the HP fold, we ignored the merged and bifurcated parts for this quantification (see Figure S9B’). We performed an additional control to account for potential effects of the temperature shift (control*^HS^*). At wL3, we find that the HP and HH folds in Np*^OXts^* are similar in width and height to those of WT and control*^HS^* (Figure 4E). This result indicates that fold deepening is not affected and that the folds do not open prematurely. At 2 hAPF, for Np*^OXts^*, the local curvature is below the detection threshold in 2 out of 3 wing discs in the region of the HP fold (Figure S9E’), consistent with a completely unfolded fold. In contrast, the HP fold is still detectable in the control*^HS^*, although it is wider than the WT (Figure 4E and Figure S9D’). The HH and HN fold of Np*^OXts^* remain folded even deeper than the WT and control*^HS^* at this stage (Figure 4E and Figure S9C). These results indicate that upon premature aECM removal, the opening of the HP fold is accelerated and the opening of the HH fold is impaired.

We next tested whether the perturbation of fold morphology in the larval wing disc prevails into the adult fly wing. As aECM has well established roles in pupal morphogenesis [19, 9], we use Np*^OXts^* and shift to an inactivated state during pupal development. We tested whether the removal of aECM before eversion, either throughout larval growth ( Np*^OXts^larva*) or during the 24h leading up to the wL3 stage ( Np*^OXts^*24*h*), results in an adult wing shape phenotype. As wing shape is both temperature- and genotype-sensitive, we used two different controls (parent Gal4 line both in-crossed and out-crossed to the Oregon-R wild type strain).

We find that the wing discs develop into relatively normal, flat wings in either perturbation (Figure 4F). However, since the perturbed folds affect mostly lateral regions of the wing disc, we hypothesized that there would be defects in the corresponding regions in the adults. Therefore, we quantified the shape and size of the adult intervein regions, which can be mapped to the different regions in the wing disc (Figure 4F, cartoon). We find that the area of the intervein regions and the wing as a whole are mostly affected by genetic background and temperature (Figure S10 and S11). In contrast, eccentricity shows a perturbation-specific effect, predominantly in two intervein regions: post L5 and L2-L3. These regions have a lower eccentricity than the genetic control and heat shock controls. This effect is more pronounced in the Np*^OXts^larva* condition than in the shorter induction of Np*^OXts^*24*h* (Figure 4F).

The L2-L3 and post-L5 regions originate from the lateral regions of the wing disc [4], which are the regions we observed to be most affected by the misshapen folds, see Figure 4C,D, Figure S8A,D, and S9B, indicating that the overexpression of Np at larval stages not only changes wing disc morphology but also affects final organ shape.

### Unfolding during wing eversion requires aECM degradation

Having established that the premature loss of aECM has an effect on fold morphology, we next address whether the physiological removal of aECM is required for unfolding of the wing disc during eversion. To do so, we prevented aECM degradation using a double knockdown of the aECM proteases Np and Stubble (Sb) (Sb*^RNAi^*; Np*^RNAi^*) [26, 24, 25].

We first investigate whether the distribution of aECM changes in the Sb*^RNAi^*; Np*^RNAi^* condition. We perform a regional knockdown in the distal region of the wing disc that affects mostly the HP fold (nub-Gal4, Figure S6) and use the Dumpy::YFP reporter to visualize aECM (Figure 5A). At 2 hAPF, we indeed observe that the aECM stays tightly connected to the apical membrane in the HP fold, whereas it detaches in the HH and HN folds (compare Figure 2C and Figure 5A). Furthermore, the HP fold remains closed, whereas the HH and HN folds appear to open normally. This difference becomes more apparent by 4 hAPF, when the HP fold remains closed and the HH fold is nearly fully open (Figure 5A and Figure S12A).

**Figure 5:**
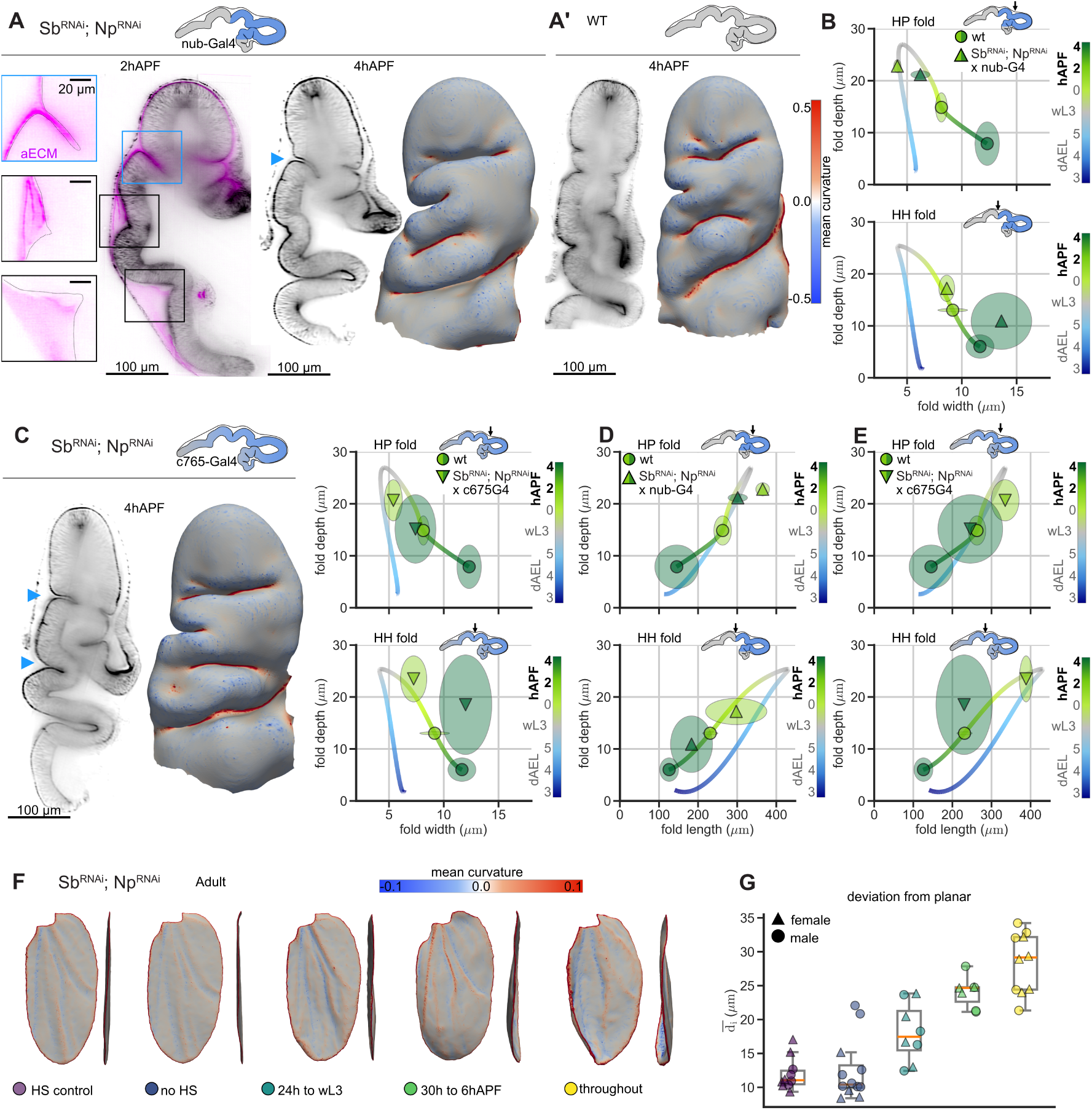
**A-E**, Cartoons indicate the expression of the Gal4 driver lines (blue) on a 2 hAPF wing disc, corresponding to the perturbation region of each experiment. **A,A’,C**, Long-axis cross-section of wing discs labeled with E-cadherin (all with Ecad::GFP, except the panel A, 2 hAPF wing disc, which is labeled with Ecad::mTomato) and head-on view of apical surface meshes (color = mean curvature *µm^−^*^1^). **A**, Sb*^RNAi^*; Np*^RNAi^* with the nub-Gal4 driver. Left: 2 hAPF wing disc of Sb*^RNAi^*; Np*^RNAi^* in a long-axis cross-section, aECM (Dumpy::YFP) in magenta and Ecad::mTomato in black. Insets: Dumpy::YFP; the apical outline of the DP is indicated by a dashed line. The positions of the insets are indicated by boxes on the cross-section. Blue box: regions covered by the nub-Gal4 driver, black: regions where the driver is not expressed. Right: 4 hAPF wing disc, the arrow indicates the HP fold that is affected by the driver. **A’**, WT 4 hAPF wing disc for comparison. **B,C,D,E**, Plots show fold shape quantification of perturbation and WT, plotted with the mean WT trajectory (line). Symbols: mean; ellipse: the extension along each axis corresponds to the 95% confidence interval of the mean. Color = developmental stage. **B**, Fold depth over width of HP and HH folds at 2 hAPF and 4 hAPF for Sb*^RNAi^*; Np*^RNAi^* in the nub-Gal4 region. **C**, Sb*^RNAi^*; Np*^RNAi^* with the c765-Gal4 driver. Left: example wing disc cross-section and head-on mesh. Right: depth over width plot. **D,E**, Fold depth over fold length for the nub-Gal4 (D) and c765-Gal4 (E) regions. **F,G**, Adult wing shape analysis for Sb*^RNAi^*; Np*^RNAi^* with the nub-Gal4 driver in different heat shock (HS) conditions. HS control: control genotype with HS 24 h prior to the wL3 stage; no HS: Sb*^RNAi^*; Np*^RNAi^* nub-Gal4*^ts^*, no HS; 24 h to wL3: Sb*^RNAi^*; Np*^RNAi^* nub-Gal4*^ts^* HS for 24 h to wL3 stage; 30 h to 6 hAPF: Sb*^RNAi^*; Np*^RNAi^* nub-Gal4*^ts^* HS for 30 h to 6 hAPF stage; throughout: Sb*^RNAi^*; Np*^RNAi^* nub-Gal4. **F**, Representative adult wing meshes colored by mean curvature (*µm^−^*^1^) in a top and lateral perspective. **G**, Quantification of adult wing deviation from planar shape for the different conditions. Color = HS conditions as indicated in F; symbol = male, female.

Fold shape quantification of 2 and 4 hAPF wing discs reveals that the HP fold in nub-Gal4; Sb*^RNAi^*; Np*^RNAi^* remains indeed closed, being deeper and narrower than in the WT (Figure 5B). We also observe that the HH fold appears sightly deeper although only significant for 2 hAPF (Figure 5B). As in WT, the HN fold remains in a bent state (Figure S12C).

To test whether the HH fold can be similarly affected by Sb*^RNAi^*; Np*^RNAi^*, we perform another knock-down, using the c765-Gal4 driver, which extends further proximally, covering the HH and, weakly, the HN fold (Figure S6B). We observe that the phenotype in the HP fold is similar to that of nub-Gal4, albeit weaker (Figure 5C and Figure S12B). In contrast, the effect on the HH fold is more pronounced than that of the nub-Gal4. This fold remains significantly deeper but becomes as wide as in the WT at 4 hAPF (Figure 5E). The HN fold is not affected by this perturbation (Figure S12D).

With both drivers, we also find that fold length is longer, in addition to being deeper (Figure 5D-E). Except for the 2 hAPF time point of the Sb*^RNAi^*; Np*^RNAi^* with nub-Gal4, all other perturbations still follow the WT trajectory, highlighting again the relationship between depth and length (see also Figure 1H).

Our results show that the presence of aECM interferes with fold opening. This finding resembles the scenario in the model where the aECM adhesion is not released and only forces beyond a minimal threshold can open the fold. To test whether this interference of fold opening by aECM has a lasting phenotype, we next address the effect on adult wing shape.

As aECM plays an important role for pupal wing morphogenesis [19, 9], and as we wished to address the effect of unfolding during wing eversion specifically, we performed a temperature-specific knockdown experiment (Sb*^RNAits^*;Np*^RNAits^*). We used the nub-Gal4 driver to focus on how HP fold unfolding affects the shape of the wing blade. We perform two controls: one testing the effect of the heat shock (HS) in a control genetic background (HS control), and the other testing potential effects of residual expression of (Sb*^RNAits^*;Np*^RNAits^*) in pupal stages (no HS). We perform three sets of perturbations: 1. Early protease knockdown 24h before wL3 stage (HS 24h to wL3); 2. Protease knockdown throughout eversion (HS 30h to 6 hAPF); 3. Protease knockdown throughout fly development (throughout). We observe that the resulting adult wings are non-planar, with the strongest phenotype occurring in the 30h to 6 hAPF and the ‘throughout’ conditions (Figure 5F,G). There is no preferred orientation of curvature (Figure S12E). As there is a clear phenotype for knockdown conditions that cease before pupal development, we conclude that the phenotype can largely be attributed to eversion stages and that unfolding during eversion is critical to attain a flat adult wing.

## Discussion

In this work, we address the question of how transient folds grow and eventually unfold to participate in the morphogenesis of the wing disc. To characterize the dynamics of fold morphology, we present a complete three-dimensional segmentation of the wing disc proper apical surface, providing an unprecedented detail of wing disc fold morphology during growth and eversion. We extend the standard approach of measuring fold shape using a cross-section with a fixed a reference point (e.g., [12, 27, 28]) by introducing a reference-free definition of fold dimensions that takes into account the full 3D fold morphology. We then identify aECM as an important component in wing disc mechanics, providing the physical link that mediates adhesion and stabilizes the folds. We use a lateral vertex model and genetic perturbations of aECM to explore its role in fold stability and unfolding. Finally, we show that perturbing fold morphogenesis in the wing disc has consequences for the adult wing shape. Perturbing the fold shape changes the proportions of the adult wing, whereas preventing unfolding during eversion leads to a curved adult wing shape. Thus, our work indicates that bilayer formation during eversion is critical for matching the dorsal and ventral wing surfaces, and errors during this period cannot be corrected later. This finding is likely due to basal connections between the two layers that form shortly after eversion and persist throughout pupal morphogenesis [23].

Our work establishes aECM as a contributor to wing disc mechanics and morphogenesis. We show that aECM provides adhesion between the apical sides of the folds. Furthermore, it likely also contributes to tissue elasticity, bending rigidity, and preferred curvature, as premature aECM removal changes the stereotypic pattern of the folds, leading to bifurcations and fold merging (Figure 4C-D, Figure S8, and S9). This result could indicate that the aECM acts as a dynamic scaffold, built by the epithelium, maintaining the fold shape as the tissue grows. The aECM may also directly affect apical cell cortex organization and mechanics. ZP-domain proteins are known to affect cytoskeletal organization, cell shape, and/or apical membrane organization in other tissues, including late pupal wings [29, 30], denticles of the larval epidermis [31], embryonic trachea [32], and the corneal lens [33]. In our TEM data, we observe aberrant apical microvilli in the Np*^OX^* condition (Figure 4B, Figure S7B,C), suggesting that aECM degradation feeds back onto microvilli formation or stability, perhaps via regulation of actin dynamics or apical membrane organization.

Interestingly, the premature removal of aECM does not lead to opening of the folds until the onset of eversion (Figure 4A-E, Figure S8, and Figure S9). This finding suggests that during eversion other forces are generated that are required for the tissue unfolding. Such forces could come from active cell dynamics in the pouch [5] and contraction of myosin cables at the periphery of the peripodial epithelium [20]. In the opposite scenario, where the aECM is not degraded at the onset of eversion, the corresponding folds remain closed (Figure 5A-E and Figure S12A,B), indicating that the aECM adhesion is strong enough to sustain the forces on the fold during eversion. Our results also suggest that aECM participates in the propagation of these eversion forces across the wing disc. For example, the opening of HH and HP folds happens at the same time in WT. In contrast, premature degradation of aECM in Np*^OX^* leads to desynchronization, with the HP opening earlier than in WT and the HH fold opening relatively later (see Figure 4D-E and Figure S9D-E’). This suggests that aECM has a role in mechanical coupling between these two folds, propagating forces between them. Furthermore, it is known that aECM in other contexts can withstand strong forces to define tissue shape during morphogenesis [19, 9, 34]. Therefore, it is plausible that it plays a similar role in the wing disc. Future experimental measurements of aECM material properties, dynamics, and composition would provide a more quantitative understanding of the mechanics of wing disc growth and eversion and its folding dynamics.

We have incorporated the aECM adhesion in the lateral vertex model to study how a fold formed in presence of aECM unfolds when an external force is applied, emulating eversion forces. We find that when simulating WT conditions, where the adhesion is removed, the unfolding trajectory parametrized by depth and width recapitulates the proportional change of fold width and depth observed in the experiments (Figure 1G, 3C, and direct comparison in Figure 3D). The opening trajectory slope is largely independent of the pulling force magnitude, indicating that such a trajectory would still be observed if the pulling forces vary over time. Interestingly, the similarity between the shape trajectories of the HP and HH folds support the model finding that opening trajectories are robust to changes in the boundary forces, as these different folds may be subject to different forces due to their position and geometry. Our model additionally predicts that unfolding is stalled when adhesion representing aECM is not removed, unless the pulling forces are sufficiently high, consistent with Sb^RNAi^ and Np^RNAi^ experiments. For higher pulling force magnitudes, the unfolding trajectory resembles ‘unzipping’, with the fold remaining very narrow. This trajectory is very similar to the time-reversed trajectory of fold growth observed in experiments, as visible in direct comparison of the two in Figure 3D, but the underlying mechanisms are likely different.

Finally, our 3D characterization of the apical surface opens the possibility to study the full wing disc architecture. In particular, our data clearly reveal that there are prominent folds on the lateral and ventral sides that are likely contributing significant material and influencing overall tissue mechanics (Figure 1B-C and Figure S1). Therefore, more work on these folds will be required to fully understand the morphogenesis of the wing. Furthermore, our quantification reveals a striking relationship between fold depth and length that remains conserved through folding and unfolding, particularly visible for the HP fold (Figure 1H). This behavior suggests that there is an underlying constraint that links these two fold dimensions. In contrast, fold depth and width are not related in the same way (Figure 1G), highlighting a possibly special relationship between length and depth. Understanding 3D wing disc morphogenesis will require a 3D model of epithelial folding that would also have to include a model of aECM mechanics and its interaction with the tissue. Having such a model would also allow us to explore whether the aECM is involved in transmitting mechanical forces between the folds during eversion, to account for the uncoupling between fold opening dynamics observed when aECM is degraded prematurely.

In conclusion, our work establishes the importance of apical ECM in fold stability and overall tissue mechanics underlying 3D tissue morphogenesis in the wing disc. More broadly, whether the fold growth and stability in other tissues is regulated using similar principles remains an interesting topic for future work.

## Acknowledgments

This work was funded with a grant to MP and NAD from the Deutsche Forschungsgemeinschaft (DFG), Project number 544201605, DY 180/2-1 and PO 3023/2-1, as well as through the Excellence program of the DFG: EXC 2068–390729961–Cluster of Excellence Physics of Life of TU Dresden, and core funding provided by the Max Planck Society to MP, FJ, JFF, and NAD. NAD additionally acknowledges Deutsche Krebshilfe (MSNZ P2 Dresden). SL’s contribution was conducted within the Max Planck School Matter to Life, supported by the German Federal Ministry of Education and Research (BMBF) in collaboration with the Max Planck Society. This work was supported by the Light Microscopy Facilities at Physics of Life, the CMCB Technology Platform at TU Dresden, and at the MPI-CBG. Authors also acknowledge assistance with Napari and the 3D apical surface extractions from Johannes Soltwedel at the BioImage Analysis Facility at Physics of Life, TU-Dresden. For assistance with the electron microscopy experiments, we are grateful to Jana Meissner at the Electron Microscopy Facility at the MPI-CBG. We also would like to acknowledge the Fly facility of the MPI-CBG, as well as Anastasia Gabrielyan and Sanika Jahagirdar in the Dye team for assistance with fly stock maintenance. Finally, we thank all the members of the Dye and Popović teams for stimulating discussions on the topic.

## Methods

Fly rearing, dissection, light sheet imaging, and processing pipeline were performed as we previously described in [35, 5] and will be explained briefly or when different from the previously published protocols.

### 1 Drosophila melanogaster preparation

#### Fly husbandry and preparation

*Drosophila melanogaster* were maintained at 25°C under 12 h light/dark cycle and fed with standard food containing cornmeal, yeast extract, soy flour, malt, agar, methyl-4-hydroxybenzoate, sugar beet syrup, and propionic acid. As WT, we used the F1 offspring of a cross between *w-;ecad::GFP*;; and *w;nub-Gal4,ecad::GFP;;*. Wing discs from male larvae or pupae at the desired stage were dissected in culture medium and immediately used for imaging or high pressure freezing.

#### Staging

Adults were flipped into a fresh food vial containing standard food and allowed to lay eggs in the morning for 3-6 h. Larvae were raised at 25°C under 12 h light/dark cycle. Imaging was performed on day 3 (74 hAEL - corresponding to the onset of fold formation), day 4 (96 hAEL), and day 5 (120 hAEL); wL3 stage wing discs were collected from upcrawling larvae with non-inflated salivary glands. Early pupa (0, 2, 4, and 6 hAPF) were staged from the time of white pupa formation.

#### Heat shock experiments

Temperature sensitive tub-Gal80^ts^ crosses were raised before and after heat shock at 18°C. Heat shocks were performed by placing the vial containing larvae/pupae in a water bath set to 30°C. To stage wL3 larvae in heat shock experiments, wL3 larvae were collected after a set time and either imaged directly or shifted back to 18°C for rearing. For prepupal stages, white prepupae were identified and marked after the set time interval and then kept in the water bath until the desired stage.

#### Drosophila melanogaster lines

Except for the UAS-Np line, all lines used in this work have been previously described and are publically available (listed in Table 1). The Np overexpression line was generated entirely by WellGenetics Inc. The Np gene was introduced into the vector pUAST-attB. To do so, the Np gene was synthesized in two parts: a 1418-bp fragment of Np N’ was synthesized with 4bp *Drosophila* Kozak sequence added to the front of the start codon; a 1774-bp fragment of Np C’ was also synthesized. A silent mutation from GAG to GAA at E671 of Np was introduced to facilitate cloning. DNA fragments were then cloned into a XhoI site of the vector pWG0773 pUAST-attB using sequence and ligation-independent cloning methods. The resulting plasmid was sequenced by Mission Biotech to verify correct cloning. Finally, the pUAST-attB-Np vector was microinjected into embryos for insertion into the ZH-86Fb site on 3R (86F8), and resulting transgenics were selected using the white marker and balanced.

**Table 1:**
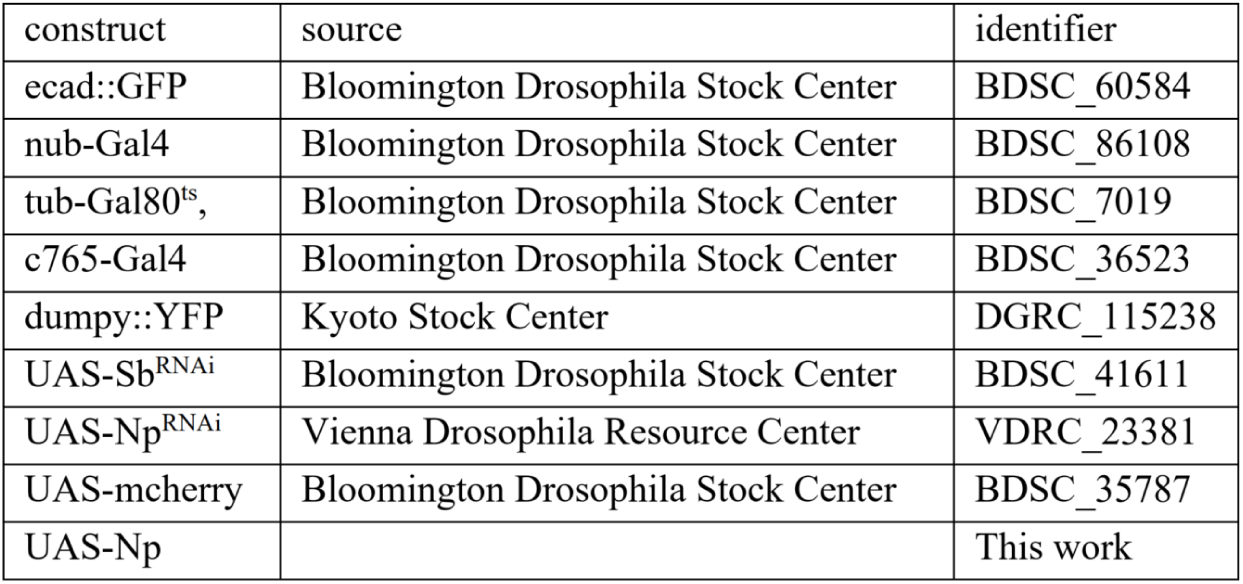
Fly strains used in this paper.

### 2 Light-sheet imaging and processing

#### Membrane labeling for live imaging

We used the FM*^T^ ^M^* 4-64 dye (Thermo Fisher Scientific, Cat T3166) for membrane labelling in a 1:1000 dilution in culture medium (Grace’s insect medium (Sigma-Aldrich, G9771, pH=6.6) and 5% Fetal bovine serum). Wing discs were incubated only for a short period (5-15 minutes) and imaged immediately afterwards to avoid labeling of intracellular vesicles by endocytosis.

#### Image acquisition and processing

Wing discs dissections and culture media were as described in [35]; prepupal dissections, imaging, and image processing was done as as described in [5]. Briefly, wing discs were dissected from larval and pupal carcasses in culture medium, and multi-angle imaging was performed on live wing discs immediately after dissection using a Zeiss Lightsheet 1 system PCO edge 4.2M Monochrome sCMOS cameras and a Plan Apo 20× [1.0 numerical aperture (NA)] water immersion objective for detection and an upgraded Zeiss Lightsheet 1 system to Z7 (Illumination: Lightsheet 7 illumination optics 10×, and detection optics 20× [1.0 NA]). Multi-angle image reconstruction was performed using BigStitcher [36].

### 3 Wing disc apical surface extraction

#### Image segmentation

Multi-angle fused image stacks were used for 3D sub-apical surface extraction. Image voxel sizes are isotropic and in the range of of 0.45 - 0.58 *µm*^3^. For each wing disc, we use Napari [37] to annotate a subset of pixels in cross-sections of the long axis of the wing disc: one in the center and 1-2 in the periphery (anterior and posterior sides). The foreground was sparsely annotated with the brush tool (pixel size = 1) on the Ecad::GFP signal on a subsection of the pouch where we had a clear signal and in 1-2 folds where the signal was slightly more blurred due to light scattering. We annotated the lateral sides as a second and the image background as a third label. We trained the Napari APOC classifier on the annotations [37, 38] with the following parameters: max depth = 2, num ensembles = 500, Features = (original gaussian blur=5, difference of gaussian=5, laplace box of gaussian blur=5) and predicted the foreground segmentation. We then used clesperanto [39] to remove small labels from this segmentation (maximum size = 100,000). We thereafter used Napari again to manually correct the segmentation in every 10th section along the AP-axis and assigned the same segmentation in five slices before and after to fill the gaps. Using this sparse segmentation, holes sometimes appear when the curvature is high in the last 30 posterior and anterior sections of the segmentation. If this was the case, additionally every 5th section in this region was corrected based on the original prediction and similarly assigned to 5 slices before and after. We then used Vedo (version ‘2023.5.0’) [40] to extract a point cloud of the segmentation and to perform sub-sampling (fraction=0.005) and smoothing (Method: moving least squares2D, f = 0.1) and open3d [41] to remove outliers (Parameters: number of neighbors=20, standard ratio = 2.0). The point cloud was converted to a density field, applying a radius of 10 pixels to close holes (using Vedo), and then to a label image (using clesperanto, connected components labeling box for all values greater than zero, followed by label erosion with a radius of 5 pixels). The Mesh was generated following the outline of the label with the “napari-process-points-and-surfaces” tool [42].

#### Mesh processing

We used open3d [41] to simplify and smooth the meshes to a target number of 100,000 triangles and denoise the mesh using an average filter with the `filter_smooth_simple` function and 5 iterations [43]. To select the sub-apical half of the mesh, we manually remove vertices of the overlaying layer using Blender [44]. We then perform hole filling using the PyTMesh `fill_small_boundaries` function (nbe=200, refine=True) [45]. Then we down-sampled the mesh with open3d by fixing, for all the meshes, a typical triangle area equal to approximately 5 *µ*m^2^, in order to mimic a typical cell area in the wing pouch region [46].

### 4 EM imaging

Wing discs were dissected in culture medium and transferred to high-pressure freezing carriers (aluminum, diameter 3 mm, depth 0.2 mm, Wohlwendt, Switzerland) in 1-Hexadecene (CH_3_(CH_2_)_13_CH=CH_2_, Merck). Fixation was performed using a Leica ICE high-pressure freezer (WT samples Figure 2A,B, S3) and a Wohlwend HPF Compact 03 ( Np*^OXts^*, Np*^OX^*, and control WT, Figure 4B, Figure S7). Freeze substitution was performed, decreasing from −90 °C to 0 °C over the time course of 50 h, in a Leica EM Automatic Freeze Substitution unit (AFS2) with 0.2% OsO4 (EMS), 3% H2O in acetone as substitution medium. Samples were washed 4x with acetone and incubated in 0.5% uranyl acetate (EMS) in acetone for 1 h at 4 °C. Samples were washed for 1 h at 4 °C and twice for 1 h at room temperature in acetone. Thereafter, acetone was gradually removed with epoxy resin (EponMethods 128 replacement, EMBed, EMS) in a 1 ml volume on a rocking table (acetone/resin 2:1 overnight; acetone/resin 1:1 for 5 h; acetone/resin 1:2 overnight; pure resin for 5 h; resin overnight; resin for 5 h). The resin was polymerized at 60°C for three days. Ultra-thin sectioning (70 nm) was performed using a Leica EM UC6 ultramicrotome on formvar-coated slot grids. Images were acquired with a FEI Tecnai 12 TEM (100kV) and an axial TVIPS CCD camera (F416, 4kx4k). Image contrast was adjusted for presentation, and tiled images were stitched using the TrakEM2 Fiji plugin [47].

### 5 Adult wing mounting, imaging, image processing

#### Dissection and mounting

Adult *Drosophila* 12-24 h post eclosion were anaesthetized with CO_2_ and fixed in isopropanol for a minimum of 5 h. One wing per fly was dissected in isopropanol using forceps. Wings were transferred to a glass cover slide. For 3D analysis of curved wings, two spacers (iSpacer(R), SuJin Lab Co. 0.2 mm deep each) were used. Wings were aligned and covered with 50% Euparal (Carl Roth, No. 7356.1) in isopropanol. After drying for 10-20 min, 75% Euparal in isopropanol was added and covered by a glass coverslip. After drying overnight, slides were sealed with nail polish.

#### Imaging

Adult wings were imaged using an automated slide scanner (Zeiss Axio Scan.Z1) of the Light Microscopy Facility, a Core Facility of the CMCB Technology Platform at TU Dresden. For 2D imaging, a Plan-Apochromat 20x/0.8 (Zeiss) objective and EDF (extended depth of focus, method wavelets) were used. For stack images, Z-stacks with 4 *µm* step-size were acquired with a Plan-Apochromat 10x/0.45 (Zeiss) objective. All images were captured with a HV-F202SCL (Hitatchi) color camera.

#### Regional analysis in flat wings

EDF images of adult wings were segmented using the Napari APOC classifier [37, 38]. A single dataset (EDF image, pixel size = 4.6 *µm*^2^ with the brush tool width = 5-10 pixel) was used as training. As foreground, the veins were annotated. The classifier was trained with the following parameters: max depth = 2, num ensembles = 500, Features = (original gaussian blur=5, difference of gaussian=5, laplace box of gaussian blur=5). The resulting segmentation was manually pruned when necessary, and the hinge region was removed. The different intervein regions were then assigned individual labels. Wing shape was analyzed for each label region using the ‘regionprops’ tool of the scikit-image library [48].

#### Curved wing analysis

Image Z-stacks of adult wings were segmented using the Napari APOC classifier [37, 38]. A single training dataset (cropped region of 1/4 of an adult wing Z-stack, pixel size = 0.88 x 0.88 x 4 *µm*) was annotated in Napari with the brush tool width = 1 pixel. As foreground, ca. 50 hair roots on the wing surface were annotated, and three different out-of-focus parts of the wing were labeled as background. The classifier was trained with the following parameters: max depth = 2, num ensembles = 500, Features = (original gaussian blur=1, difference of gaussian=1, laplace box of gaussian blur=1) [37, 38]. The Hinge region was then manually labeled in Napari and cropped from the prediction. We then generated a point cloud from the segmentation using Vedo [40] and performed sub-sampling, outlier removal, and smoothing. If, upon inspection, the point cloud had large gaps, we filled these regions manually in Napari. Thereafter, using Vedo, the point cloud was transformed into a density image with radius=15 and all non-zero values classified as foreground. The resulting binary was eroded once using skimage.morphology with an isotropic footprint of 10 [48]. We then used Napari for mesh creation using the “napari-process-points-and-surfaces” tool [42]. We then reduced the number of triangles to 100,000 (open3d simplify-quadric-decimation) and smoothed the surface with 5 iterations using open3d (filter-smooth-simple)[41]. The visualizations were generated using Paraview [49], coloring mesh vertices using their mean curvature (see Methods, section 6).

#### Plane fit derivation

To calculate the deviation from a plane, we compute the covariance matrix of the mesh vertices and identify the best-fit plane as the one whose normal corresponds to the direction of least variance in the vertex positions. We then calculate the perpendicular distance from each vertex to this plane and report the average distance per wing. To analyze the curvature of these wings, we first split each wing mesh into a dorsal and ventral component. This is achieved by cutting the mesh at the wing margin, as identified by a mean curvature threshold of 0.15 *µm^−^*^1^. Thereafter, the area weighted average mean curvature was computed for the larger of the two components. The sign of mean curvature was chosen by orienting the mesh surface normal vectors towards the dorsal side (Figure S12E).

### 6 Mesh curvature

All surfaces were analyzed as triangular meshes. To study the fold regions, we employ the mean curvature calculated at the vertex level using the discrete Laplace-Beltrami operator computed using the “cotan” formula [50]:

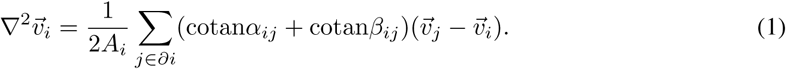

Here, *α_ij_* and *β_ij_* are the angles opposite to the edge *ij* in the two triangles that share the edge *ij*. *A_i_* is the Voronoi-cell area associated with the vertex *i*, computed as 1*/*3 of the sum of areas of the triangles adjacent to the vertex *i*. The magnitude of the mean curvature of the vertex *i* is obtained as:

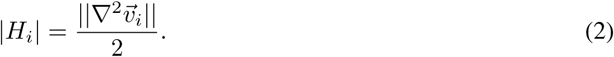

The sign of the mean curvature is defined by our choice of the mesh surface orientation, where we choose the normal vector of the mesh surface to point in the direction away from the wing disc tissue.

### 7 Fold regions segmentation

Each fold region is obtained by a clustering method that considers both mean curvature and spatial proximity. First, each vertex is flagged as a candidate vertex to belong in a fold region by simple thresholding on its mean curvature *H*_min_ *≤ H_i_ ≤ H*_max_. Then, considering all such vertices, we cluster them spatially by euclidean distance using a threshold *d*. Finally, we inflate the obtained clusters by including all vertices at a distance *d^′^* from any of the vertices in the region. While precise values for the parameters depend on the sample and stage, they remain on the order of *H*_min_ = 0.2, *H*_max_ = 3, *d* = 5, and *d^′^* = 15.

### 8 Bottom curve identification

The bottom curve of a fold is defined via an optimization procedure. First, in each fold region we identify two endpoint subregions *E*_1_ and *E*_2_. To do so, we first project the vertex coordinates along the longest direction of variation of the fold coordinates, by means of a simple Principal Component Analysis, thus obtaining projection coordinates *{ζ_i_}*. We take the vertices in the first endpoint subregion *E*_1_ as those with ζ*_i_ ≤* ζ_0.05_, where ζ_0.05_ is the 5% quantile of the coordinates and in endpoint subregion *E*_2_ as those with ζ*_i_* > ζ_0.95_, the 95% quantile. Then, we construct the adjacency graph of vertices belonging to the fold *G*. For each pair of vertices (*v_rmstart_, v_rmend_*) such that *v*_start_ *∈ E*_1_ and *v*_end_ *∈ E*_2_, we find the optimal path in *G* that connects them and minimizes the cost function:

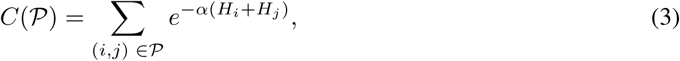

where (*i*, *j*) are the edges belonging to the candidate path, *H_i_* the mean curvature, and *α* = 2. This cost function promotes paths with overall positive mean curvature. We thus obtain a collection of paths *P_1_, …, P_M_* and merge them by applying the Dynamic Time Warping (DTW) algorithm directly on the vertex coordinates. Such algorithm outputs an ordered collection of points *b⃗*_1_, …, *b⃗_T_*. To each point *b⃗_t_* we obtained also an alignment error *δ_t_*, directly from the alignment algorithm, which measures how much the original paths are spread around the aligned point *b⃗_t_*. We use this error to discard badly-aligned points via a percentile-based elimination, where we discard the top 20% of worst aligned points, via an empirical validation. Finally, we use a spline to fit these points, which we define to be the bottom fold curve *b⃗*(*t*).

### 9 Cross-sections computation

Having obtained the fold bottom curve *b⃗*(*t*) = (*b_x_*(*t*)*, b_y_*(*t*)*, b_z_*(*t*)), we now define a set of cross-sections along the curve, equally spaced by approximately *λ* = 4.3 *−* 4.8 µm depending on the developmental stage. Each cross-section of the fold at point *b⃗*(*t*) contains a set of mesh vertices *V_t_*, chosen from the fold region, whose distance from the cross-section plane is smaller than *λ*. The points contained in *V_t_* span a region much larger than the fold region and thus need to be restricted. We do so by the following procedure.

First, we define an initial reference frame on the cross-section plane. We construct the reference frame by exploiting the mesh surface normal vectors. We look at vertices in the mesh within a distance *d^∗^* = *µm* from the fold bottom, and we average corresponding normal vectors. We then project this average to the cross-section plane and normalize it to obtain vector *N*^^^(*t*), which defines the *y*-axis of the initial reference frame. Finally, the *x*-axis is defined by the vector *B*^^^(*t*), obtained as the cross product between *T*^^^(*t*) and *N*^^^(*t*). We now express positions of each point in *V_t_* using the coordinates *x_i_* and *y_i_* of the initial reference frame. Moreover, as each point in *V_t_* has an associated normal vector in the mesh, we project these normal vectors to the cross-section plane, which after normalization read *n*^*_i_* = (cos *θ_i_,* sin *θ_i_*) in the reference frame.

We then create the relevant subset of points in *V_t_* in two steps. First, we discard the points for which sin *θ_i_* is too negative (in a stage-dependent manner, limiting thus angles, typically, such that sin *θ_i_ ≥ −*0.3). Second, we use an adaptive method, starting from a small subregion of points satisfying *|x_i_| ≤ x*_max_ and also for *y*_max_ such that 0 *≤ y_i_ ≤ y*_max_. The initial *x*_max_ goes from 10 *µm* to 20 *µm* as we go through stages, and similarly *y*_max_ from 24 *µm* to 48 µm. We then increase both *x*_max_ and *y*_max_ by a factor *f* = 0.01, and check whether the included maximum *y* coordinates saturate past a horizontal sufficient distance from the bottom (taken as 10% of the total horizontal span). If the condition is fulfilled, the procedure is finished. If not, we repeat the procedure. This choice is driven by the notion that tissue in the fold is invaginated, while it flattens out outside of the fold. Once this adaptive procedure stops, we are left with a final set *V_t_ ^′^* of mesh vertices projections on the cross-section plane that we denote fold points.

### 10 Fold height and width computation

To define fold height and width, we first consider the shape traced by fold points in each cross-section. Since these shapes can be asymmetrical, we separately analyse each side of the cross-section, separated by the bottom curve point. In this way, we obtain ‘left’ and ‘right’ subsets of the fold points *i* by criterion *x_i_* < 0 and *x_i_* > 0, respectively. For each side of the section, we construct the upper envelope as:

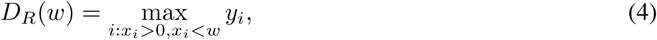

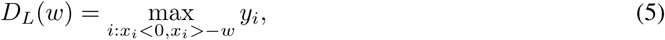

where *w* is a coordinate of each envelope along *x*-axis. We define left and right fold depth *D_L_* and *D_R_* via integral averaging procedure.

Namely, we first compute left and right widths *W_L_* and *W_R_* as integral averages of the profile:

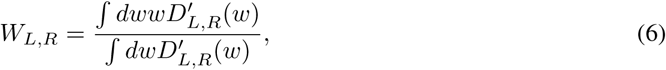

where the prime symbol denotes differentiation. As the envelope is a step-wise function, we do discrete differentiation and integration via the Python Numpy library “np.gradient” and “np.trapz” functions. The left and right depths *D_L_* and *D_R_* are thus obtained by finding the position of the envelopes at *W_L_* and *W_R_* via linear interpolation. Using the same procedure, we also compute the variance associated to this measurement:

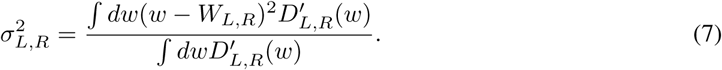

We use the variance to weight each measurement to obtain a single, per cross-section, measurement as

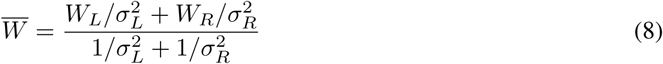

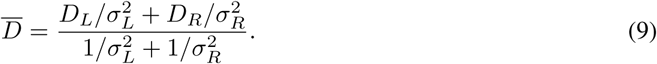

Finally, the fold depth *D* and width *L* are the integral averages of all the cross-sections along the fold bottom.

### 11 Measurement of fold shape

To represent developmental dynamics of fold depth, width, and length in the wild type, we construct the shape trajectory from the measurements. For this, we consider at each developmental stage a single replicate and fit a cubic spline. We repeat this for all possible combinations of replicates in each stages (see Table 2) to obtain an ensemble of spline curves. We interpolated each spline at 200 equally spaced time points and determine the average spline curve over the ensemble of spline curves, shown by solid thick line all figures. We estimate the uncertainty interval at each of the time points as the standard deviation over the same ensemble, shown in by error bars in Figures 1E,G,H and Figure 3D.

**Table 2:**
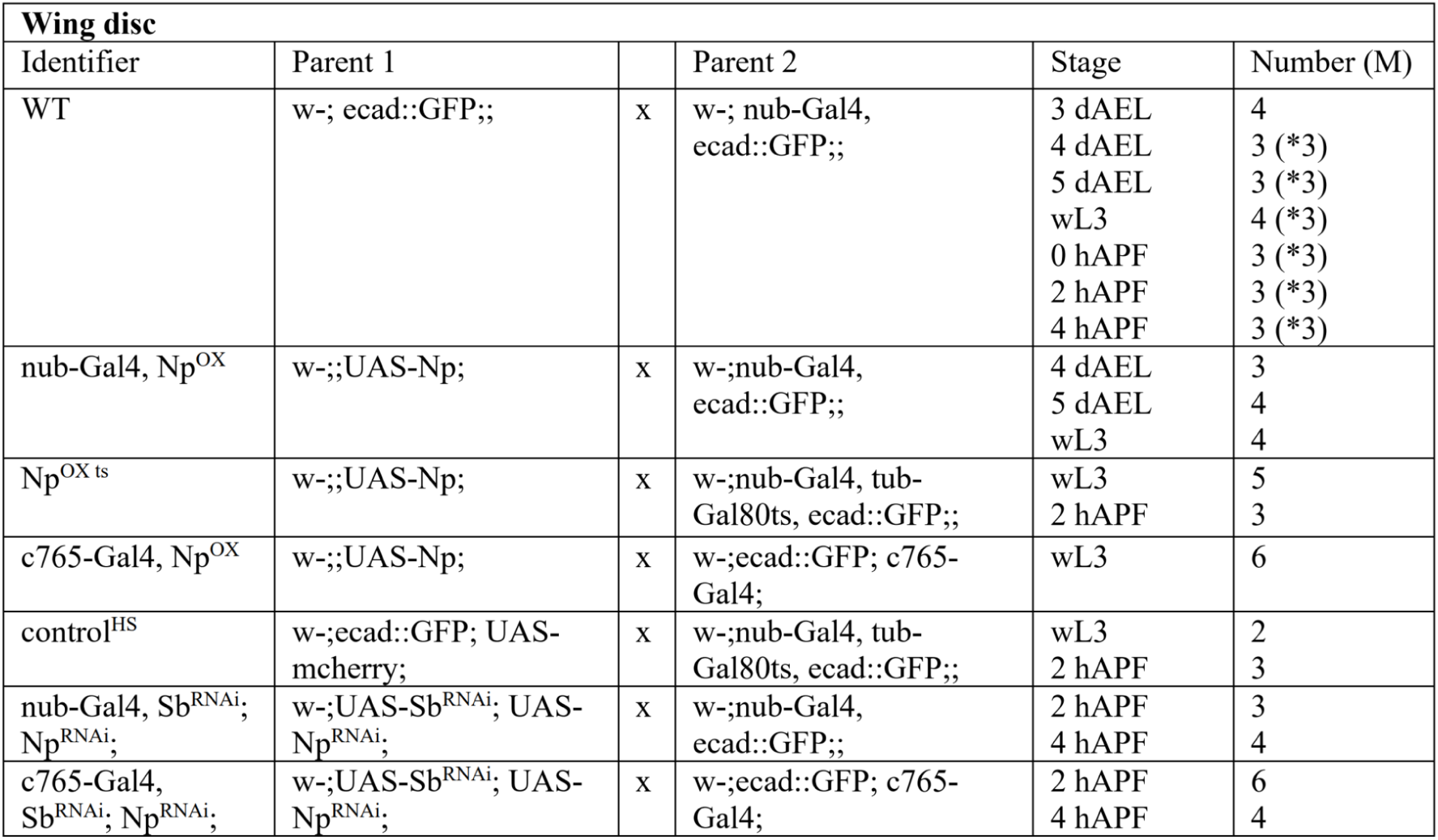
Wing discs analyzed in this work; The numbers in brackets (*) indicate how many of these samples have been previously published in [5]; Only male (M) wing discs were analyzed.

**Table 3:**
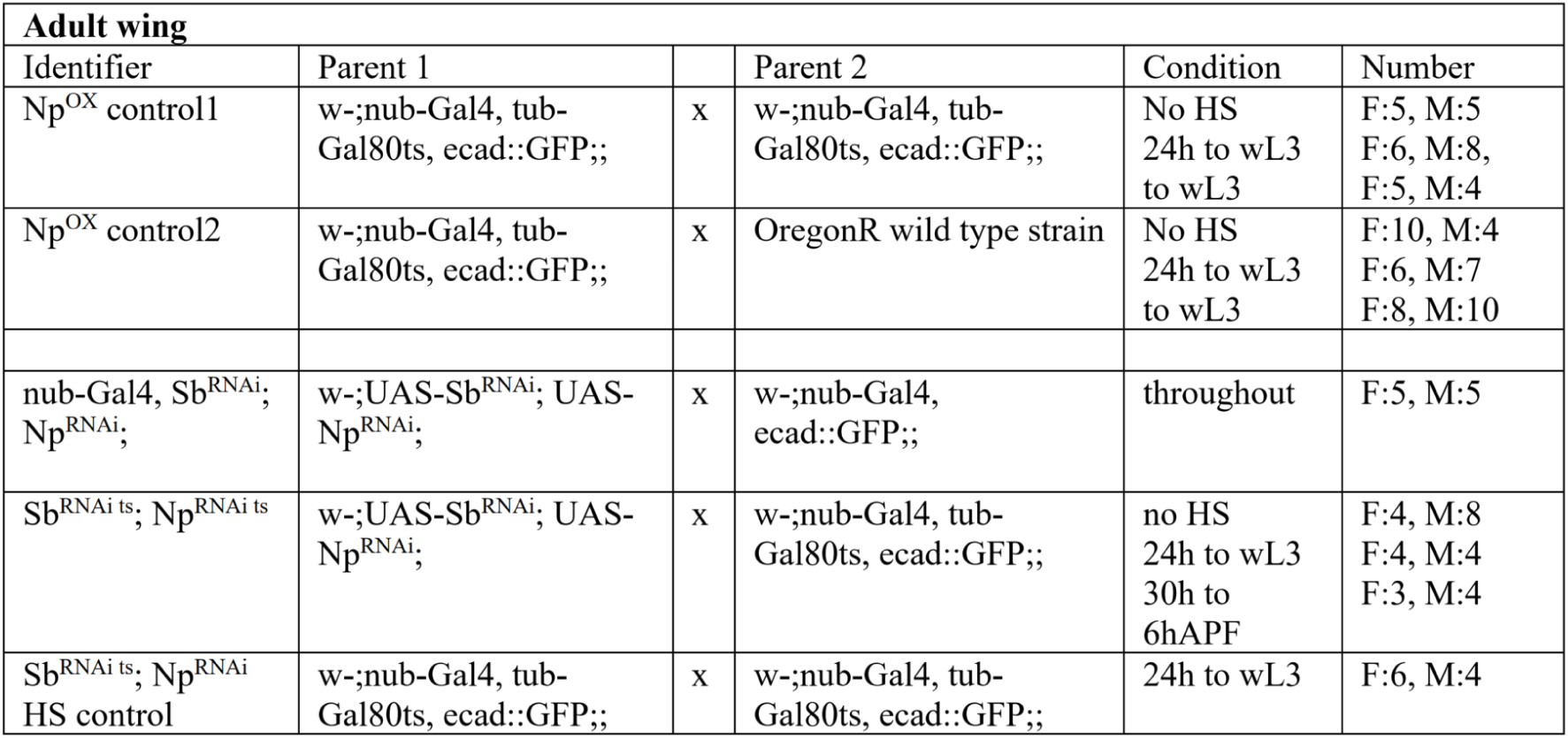
Adult wing data; male = M, female = F.

When comparing measurements of the fold dimensions between the WT and genetic perturbations in Figures 4E, 5B-E, S8D’, D”, E’, and S9C, we estimate the uncertainty interval at each developmental stage using 1000 independent bootstrap samples of our data. We report the statistic mean and the 95% confidence interval of the bootstrap samples as the data point and an ellipse around the mean.

