## Supplemental Material for "Apical extracellular matrix regulates fold morphogenesis in the *Drosophila* wing disc"

### 1 Lateral vertex model

#### 1.1 Tissue model

We denote the $N_v$ vertices of the apical surface as $\vec{r}_{a,i} = (x_{a,i}, y_{a,i})_{i=1}^{N_v}$ and of the basal surface as $\vec{r}_{b,i} = (x_{b,i}, y_{b,i})_{i=1}^{N_v}$. Each of the $N = N_v - 1$ cells is indexed by $c = 1 \dots N$. The lengths of the apical and basal surfaces for each cell reads:

$$\ell_{a,c} = ||\vec{r}_{a,c+1} - \vec{r}_{a,c}|| \tag{10}$$

$$\ell_{b,c} = ||\vec{r}_{b,c+1} - \vec{r}_{b,c}|| \tag{11}$$

while the length of each lateral surface:

$$\ell_{l,i} = ||\vec{r}_{a,i} - \vec{r}_{b,i}|| \tag{12}$$

Finally the area of a cell is computed using the shoelace formula:

$$A_c = \frac{1}{2}||\vec{r}_{b,c} \times \vec{r}_{b,c+1} + \vec{r}_{b,c+1} \times \vec{r}_{a,c+1} + \vec{r}_{a,c+1} \times \vec{r}_{a,c} + \vec{r}_{a,c} \times \vec{r}_{b,c}|| \tag{13}$$

The energy function of the model reads then:

$$E = \sum_c \frac{K}{2}(A_c - A_{0,c})^2 + \Gamma_{a,c}\ell_{a,c} + \Gamma_{b,c}\ell_{b,c} + \sum_i \Gamma_{l,i}\ell_{l,i} - F(x_{a,N_v} - x_{a,0}) - F(x_{b,N_v} - x_{b,0}) \tag{14}$$

where the last term appears as we apply a compressive $F = F_{\text{push}} < 0$ or extensile force $F = F_{\text{pull}} > 0$.

#### 1.2 Adhesion

The total adhesion energy between two 1D adhesive surfaces is given by a double integral of a potential function over the two surfaces:

$$U = \int\int d\vec{r}_1 d\vec{r}_2 V(|\vec{r}_1 - \vec{r}_2|), \tag{15}$$

where position vectors $\vec{r}_{\{1,2\}}$. $V$ is a pairwise interaction between two infinitesimal elements on each surface. Here, a generalized Lennard-Jones potential is used to simulate long-range attraction and short-range repulsion, which only depends on the distance between the two elements, $r = |\vec{r}_1 - \vec{r}_2|$:

$$V(|\vec{r}_1 - \vec{r}_2|) = V(r) = \gamma\big[\big(\frac{r_{\text{min}}}{r}\big)^{2n} - 2\big(\frac{r_{\text{min}}}{r}\big)^{n}\big], \tag{16}$$

where $\gamma$ characterizes the adhesion strength and $r_{\text{min}}$ determines the equilibrium distance between two bodies. In this generalized Lennard-Jones potential, the exponents for adhesive and repulsive interactions are allowed to vary, where $n = 6$ corresponds to the original Lennard-Jones potential. In this study, a smaller exponent, $n = 4$ is used to reduce the complexity of the analytic expression for $U$.

To simplify the calculation, a system of two adhesive surfaces is rotated such that one of the surfaces lies horizontally along the $x$-axis. This horizontal surface, denoted as surface 2, is defined by two endpoints, $(0, 0)$ and $(L_*, 0)$, where $L_*$ specifies the length. The other surface, denoted as surface 1, has other two endpoints, $(x_L, y_L)$ and $(x_R, y_R)$. Then the double integral, Eq. 15, becomes

$$\begin{aligned} U &= \int d\vec{r}_1 \int d\vec{r}_2 V(\vec{r}_1, \vec{r}_2) \\ &= \int d\vec{r}_1 \Phi(\vec{r}_1) \\ &= L \int_0^1 ds \Phi(\vec{r}_s). \end{aligned} \tag{17}$$

From the first to the second line, a function $\Phi$ is introduced as a single integral of a potential $V$ over surface 2. This function depends on the position vector $\vec{r}_1$ on surface 1. From the second to the third line, surface 1 is parameterized as $\vec{r}_s = (x_s, y_s)$ with $x_s = (1-s)x_L + sx_R$, $y_s = (1-s)y_L + sy_R$ and $s$ spanning from 0 to 1. $L = \sqrt{(x_R - x_L)^2 + (y_R - y_L)^2}$ denotes the length of this surface. The pairwise interaction $V$ involves two power functions of $1/r$ with the exponents of 8 for repulsive and 4 for adhesive potential. The integration of each power law $(1/r)^k$ can be defined as

$$A_k = L \int_0^1 ds B_k(\vec{r}_s), \tag{18}$$

where

$$B_k(\vec{r}_s) = \int_0^{L_*} dl \frac{1}{[(x_s - l)^2 + y_s^2]^{k/2}}, \tag{19}$$

for $k = 4$ and 8. Note that $B_k$ depends on $x_s$, $y_s$ and $L_*$, and $A_k$ depends on $x_L$, $y_L$, $x_R$, $y_R$ and $L_*$.

The total energy can be expressed as

$$U = \gamma(A_8 \cdot r_{\text{min}}^8 - 2A_4 \cdot r_{\text{min}}^4), \tag{20}$$

where

$$B_4(\vec{r} = (x, y)) = \frac{1}{2y^3} \left( \frac{(L_* - x)\, y}{(L_* - x)^2 + y^2} + \tan^{-1}\left( \frac{L_* - x}{y} \right) \right) \tag{21}$$

$$\begin{aligned} B_8(\vec{r} = (x, y)) =& \frac{1}{48y^7} \left( (L_* - x) \left( \frac{15y}{(L_* - x)^2 + y^2} + \frac{10y^3}{((L_* - x)^2 + y^2)^2} \right.\right. \\ & \left.\left. + \frac{8y^5}{((L_* - x)^2 + y^2)^3} \right) + 15 \tan^{-1}\left( \frac{L_* - x}{y} \right) \right) \end{aligned} \tag{22}$$

$$
\begin{aligned}
A_4/L =& \frac{1}{4}\left(\frac{L_*\left(y_L-y_R\right)^2}{y_L y_R\left(y_L x_R-x_L y_R\right)\left(-y_L x_R+x_L y_R+L_*\left(y_L-y_R\right)\right)}\right.\\
&+\frac{1}{y_L-y_R}\left(\left(\left(x_L-x_R\right)^2+\left(y_L-y_R\right)^2\right)\right.\\
&\left(\frac{\tan^{-1}\left(\frac{-x_L x_R-y_L y_R+x_R^2+y_R^2}{x_L y_R-y_L x_R}\right)-\tan^{-1}\left(\frac{-x_L x_R-y_L y_R+x_L^2+y_L^2}{y_L x_R-x_L y_R}\right)}{\left(y_L x_R-x_L y_R\right)^2}\right.\\
&\left.+\frac{\tan^{-1}\left(\frac{-x_L x_R+L_*\left(x_R-x_L\right)-y_L y_R+x_L^2+y_L^2}{y_L x_R-x_L y_R-L_*\left(y_L-y_R\right)}\right)-\tan^{-1}\left(\frac{-x_L x_R+L_*\left(x_L-x_R\right)-y_L y_R+x_R^2+y_R^2}{-y_L x_R+x_L y_R+L_*\left(y_L-y_R\right)}\right)}{\left(-y_L x_R+x_L y_R+L_*\left(y_L-y_R\right)\right)^2}\right)\\
&\left.\left.+\frac{\tan^{-1}\left(\frac{L_*-x_R}{y_R}\right)+\tan^{-1}\left(\frac{x_R}{y_R}\right)}{y_R^2}-\frac{\tan^{-1}\left(\frac{x_L}{y_L}\right)+\tan^{-1}\left(\frac{L_*-x_L}{y_L}\right)}{y_L^2}\right)\right)
\end{aligned}
\tag{23}
$$

$$
\begin{aligned}
A_8/L =& \frac{1}{288}\left(\frac{3}{y_L y_R}\left(5\left(x_L^4+x_R^4\right)-20 x_L x_R\left(x_L^2+x_R^2\right)-10\left(x_L-x_R\right)^2\left(y_L-y_R\right)^2\right.\right.\\
&\left.+30 x_L^2 x_R^2+\left(y_L-y_R\right)^4\right)\left(y_L-y_R\right)\left(\frac{1}{\left(x_R y_L-x_L y_R\right)^5}+\frac{1}{\left(-x_R y_L+L_*\left(y_L-y_R\right)+x_L y_R\right)^5}\right)\\
&-\frac{15}{y_L^2 y_R^2}\left(x_L-x_R\right)\left(\left(x_L-x_R\right)^2-\left(y_L-y_R\right)^2\right)\left(y_L^2-y_R^2\right)\left(\frac{1}{\left(x_R y_L-x_L y_R\right)^4}\right.\\
&\left.-\frac{1}{\left(-x_R y_L+L_*\left(y_L-y_R\right)+x_L y_R\right)^4}\right)+\frac{5}{y_L^3 y_R^3}\left(3\left(x_L-x_R\right)^2-\left(y_L-y_R\right)^2\right)\\
&\left(y_L^3-y_R^3\right)\left(\frac{1}{\left(x_R y_L-x_L y_R\right)^3}+\frac{1}{\left(-x_R y_L+L_*\left(y_L-y_R\right)+x_L y_R\right)^3}\right)\\
&-\frac{15}{y_L^4 y_R^4}\left(x_L-x_R\right)\left(y_L^4-y_R^4\right)\left(\frac{1}{\left(x_R y_L-x_L y_R\right)^2}-\frac{1}{\left(-x_R y_L+L_*\left(y_L-y_R\right)+x_L y_R\right)^2}\right)\\
&+\frac{15}{y_L^5 y_R^5}\left(y_L^5-y_R^5\right)\left(\frac{1}{x_R y_L-x_L y_R}+\frac{1}{-x_R y_L+L_*\left(y_L-y_R\right)+x_L y_R}\right)\\
&+\frac{12}{\left(y_L x_R-x_L y_R\right)^3}\left(\frac{2 x_L y_R\left(x_L-x_R\right)-y_L\left(\left(y_L-y_R\right)^2+x_L^2-x_R^2\right)}{\left(x_L^2+y_L^2\right)^2}\right.\\
&\left.+\frac{2 x_R y_L\left(x_L-x_R\right)+y_R\left(\left(y_L-y_R\right)^2-x_L^2+x_R^2\right)}{\left(x_R^2+y_R^2\right)^2}\right)+\frac{12}{\left(-x_R y_L+x_L y_R+L_*\left(y_L-y_R\right)\right)^3}\\
&\left(\frac{2 x_R y_L\left(x_L-x_R\right)+y_R\left(\left(y_L-y_R\right)^2-x_L^2+x_R^2\right)-2 L_*\left(x_L-x_R\right)\left(y_L-y_R\right)}{\left(\left(x_R-L_*\right)^2+y_R^2\right)^2}\right.\\
&\left.+\frac{2 x_L y_R\left(x_L-x_R\right)-y_L\left(\left(y_L-y_R\right)^2+x_L^2-x_R^2\right)+2 L_*\left(x_L-x_R\right)\left(y_L-y_R\right)}{\left(\left(x_L-L_*\right)^2+y_L^2\right)^2}\right)\\
&+\frac{6}{\left(x_R y_L-x_L y_R\right)^5}\left(-\frac{1}{x_L^2+y_L^2}\left(\left(x_L-x_R\right)\left(y_L-y_R\right)^2\left(-13 x_R y_L+12 x_L y_L+x_L y_R\right)\right.\right.\\
&+2 y_L\left(y_L-y_R\right)^4+5 x_L^4\left(2 y_L-3 y_R\right)-5 x_L^3 x_R\left(5 y_L-9 y_R\right)+15 x_L^2 x_R^2\left(y_L-3 y_R\right)\\
&\left.+5 x_L x_R^3\left(y_L+3 y_R\right)-5 x_R^4 y_L\right)+\frac{1}{x_R^2+y_R^2}\left(\left(x_L-x_R\right)\left(y_L-y_R\right)^2\left(13 x_L y_R-12 x_R y_R-x_R y_L\right)\right.\\
&+2 y_R\left(y_L-y_R\right)^4-5 x_R^4\left(3 y_L-2 y_R\right)+5 x_L x_R^3\left(9 y_L-5 y_R\right)+15 x_L^2 x_R^2\left(-3 y_L+y_R\right)\\
&\left.\left.+5 x_L^3 x_R\left(3 y_L+y_R\right)-5 x_L^4 y_R\right)\right)+\frac{6}{\left(-x_R y_L+L_*\left(y_L-y_R\right)+x_L y_R\right)^5}
\end{aligned}
$$

$$\left(\frac{1}{(L_* - x_L)^2 + y_L^2}\left((x_L - x_R)(y_L - y_R)^2 (13x_R y_L - 12x_L y_L - x_L y_R) - 2(y_L - y_R)^4 y_L\right.\right.$$
$$+5x_R^4 y_L + 5x_L^3 x_R (5y_L - 9y_R) - 15x_L^2 x_R^2 (y_L - 3y_R) - 5x_L x_R^3 (y_L + 3y_R) - 5x_L^4 (2y_L - 3y_R)$$
$$\left.+L_* (x_L - x_R)(y_L - y_R)\left(15(x_L - x_R)^2 - (y_L - y_R)^2\right)\right) - \frac{1}{(L_* - x_R)^2 + y_R^2}$$
$$\left((x_L - x_R)(y_L - y_R)^2 (-13x_L y_R + 12x_R y_R + x_R y_L) - 2(y_L - y_R)^4 y_R + 5x_L^4 y_R\right.$$
$$-5x_L^3 x_R (3y_L + y_R) + 15x_L^2 x_R^2 (3y_L - y_R) - 5x_L x_R^3 (9y_L - 5y_R) + 5x_R^4 (3y_L - 2y_R)$$
$$\left.\left.+L_* (x_L - x_R)(y_L - y_R)\left(15(x_L - x_R)^2 - (y_L - y_R)^2\right)\right)\right) + \frac{15}{y_L - y_R}$$
$$\left(\left((x_L - x_R)^2 + (y_L - y_R)^2\right)^3 \left(\frac{\tan^{-1}\left(\frac{-x_L x_R - y_L y_R + x_R^2 + y_R^2}{-x_R y_L + x_L y_R}\right) - \tan^{-1}\left(\frac{-x_L x_R - y_L y_R + x_L^2 + y_L^2}{x_R y_L - x_L y_R}\right)}{(x_R y_L - x_L y_R)^6}\right.\right.$$
$$\left.+\frac{\tan^{-1}\left(\frac{-x_L x_R + L_*(x_R - x_L) - y_L y_R + x_L^2 + y_L^2}{x_R y_L - L_*(y_L - y_R) - x_L y_R}\right) - \tan^{-1}\left(\frac{-x_L x_R - y_L y_R + L_*(x_L - x_R) + x_R^2 + y_R^2}{-x_R y_L + L_*(y_L - y_R) + x_L y_R}\right)}{(-x_R y_L + L_*(y_L - y_R) + x_L y_R)^6}\right)$$
$$\left.\left.+\frac{\tan^{-1}\left(\frac{L_* - x_R}{y_R}\right) + \tan^{-1}\left(\frac{x_R}{y_R}\right)}{y_R^6} - \frac{\tan^{-1}\left(\frac{L_* - x_L}{y_L}\right) + \tan^{-1}\left(\frac{x_L}{y_L}\right)}{y_L^6}\right)\right) \tag{24}$$

#### 1.3 Fold preparation

Simulations of the lateral vertex model were carried out by energy minimization via a simple gradient descent. To choose the initial parameters, we set ourselves in a free tissue case. We then can compute the flat and free tissue equilibrium geometry by minimizing the single cell energy assuming $\Gamma_a = \Gamma_b = \Gamma$ and thus a rectangular equilibrium shape $\ell_a = \ell_b = w$ and $\ell_l = h$:

$$E_{\text{cell}} = 2\Gamma w + \Gamma_\ell h + \frac{K}{2}(wh - A_0)^2 \tag{25}$$

The condition for minimization imply:

$$2\Gamma + K(wh - A_0)h = 0 \tag{26}$$

$$\Gamma_l + K(wh - A_0)w = 0 \tag{27}$$

which in turn can be recast to:

$$K(wh - A_0) = -\frac{2\Gamma}{h} \implies \Gamma_l = \frac{2\Gamma w}{h} \tag{28}$$

The formula above fixes the initial lateral tension. By eliminating now $\Gamma_l$ we arrive to:

$$\frac{2\Gamma}{h} = K(A_0 - wh) \implies A_0 = wh + \frac{2\Gamma}{hK} \tag{29}$$

So we choose $A_0$ and $\Gamma_l$ to have equilibrium values $w = w_0 = 1$ and $h = h_0 = 16$, where we set $K = 0.5$. With these choices, the corresponding unit of force reads $F_0 = 2Kw_0^3 = 1$.

For the fold preparation, we choose an initial tissue of $N_c = 201$ cells (see Figure S4). To trigger fold formation, we start from a flat tissue and we lower the basal tension by 40% and increase the lateral one by 100% in a region of $N_f = 9$ cells around the central cell. We then use a fixed pushing force applied at the tissue boundary of $F_{\text{push}} = 1$ and vary the adhesion strength $\gamma$. The change in tension at the center coupled with the boundary force induces an invagination of the tissue, which is stabilized by the adhesive forces. Once the fold has formed, we remove cells outside the fold, keeping a region of end-to-end length $L \sim 40w_0$, corresponding to around 120 cells. After this, we keep the new cut tissue at constant boundary displacement, and we find the force $F_i$ that would keep the fold stable at equilibrium.

#### 1.4 Unfolding simulations

Once we have the initial folded tissue (see Figure S4), we apply a variable boundary force $F_{\text{pull}}$ to simulate extensile stress. We moreover modify the boundary conditions to maintain apical and basal boundary vertices at the same horizontal velocity. To reach $F_{\text{pull}}$, we start from $F_i$ and gradually bring it to $F_{\text{pull}}$. This is done to avoid sudden force changes. The two different scenarios correspond to either keeping the original adhesion strength $\gamma$ or setting it to zero $\gamma = 0$. The energy minimization is then executed to observe either a complete unfolding when no adhesion is present or when the energy plateaus without tissue unfolding.

#### 1.5 Fold shape measurements and comparison with experimental data

To measure the width and depth trajectory in the simulations, as shown in Figure 3C, we adopt the method developed to quantify fold cross-section width and height from experimental data, and apply it to the apical side of the simulation cells.

When comparing the simulation results with experimental data in Figure 3D, we take into account the fact that the segmentation of the surface is sub-apical (see Figure 1A and Figure S1B). Therefore, to compare the simulation results with experimental data, instead of using the apical surface of model cells we extract a line corresponding to an offset $M$ below the actual surface. We map the model units to $\mu m$ by matching the initial fold depth in the model to the HP and HH fold depth at wL3 stage experiments. In this procedure we select $M = 2.5\ \mu m$.

### 2 Supplement Figures and Data

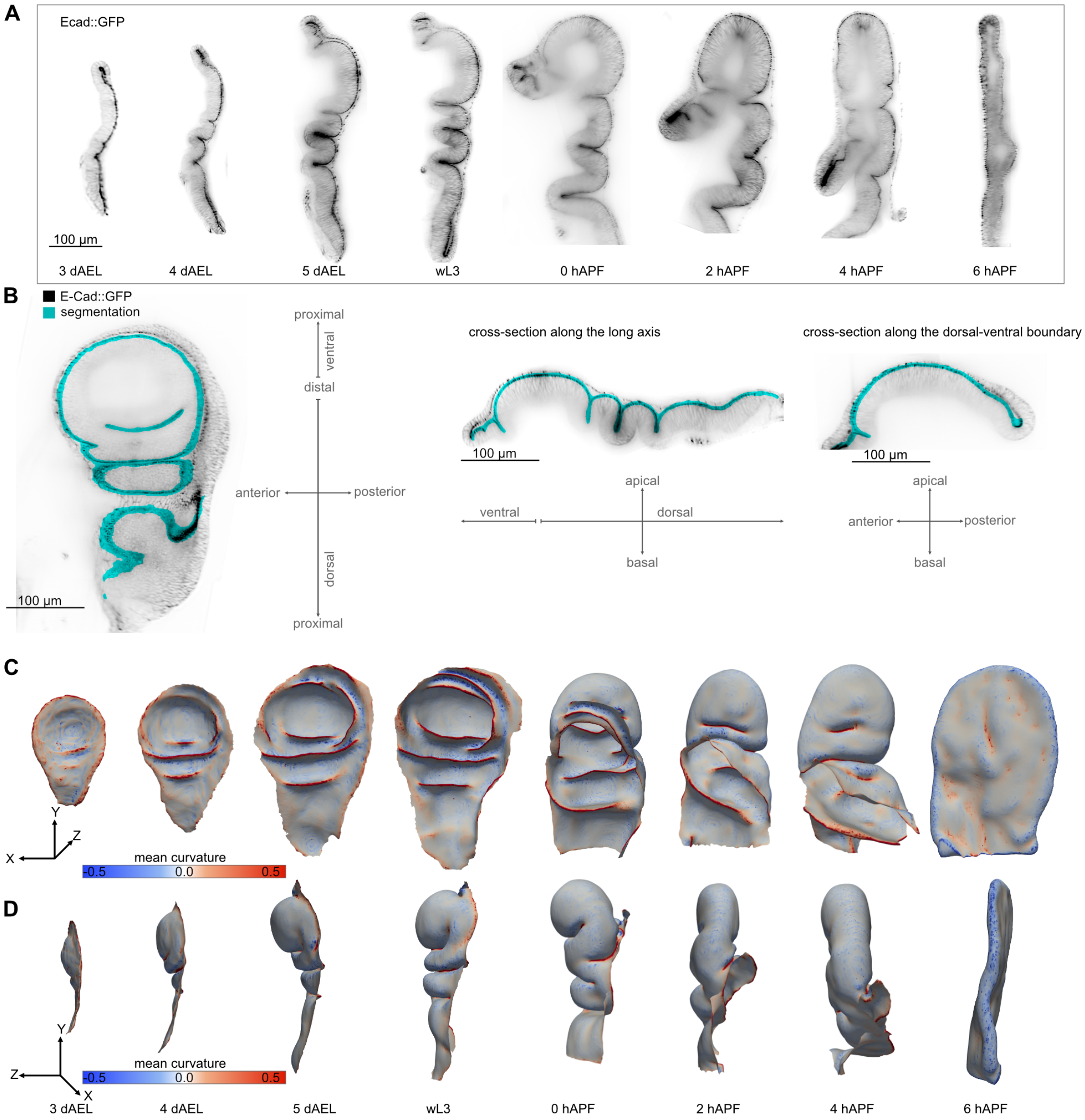


Figure S1: **A**, Long-axis cross-section of the wing disc over developmental time. Label = Ecad::GFP. **B**, Apical segmentation on an exemplary wL3 wing disc; different views are shown. Left: slice in the apical basal axis. Middle: long-axis cross-section. Right: cross-section along the dorsal-ventral boundary. Anterior is left for the left and right images; ventral is left for the middle image. **C, D**, Apical surface meshes of the wing discs in panel A, and Figure 1 A, rotated to show a different view. Color represents mean curvature ($\mu m^{-1}$). **C**: View from the basal side; anterior is right; **D**: View from the posterior side.

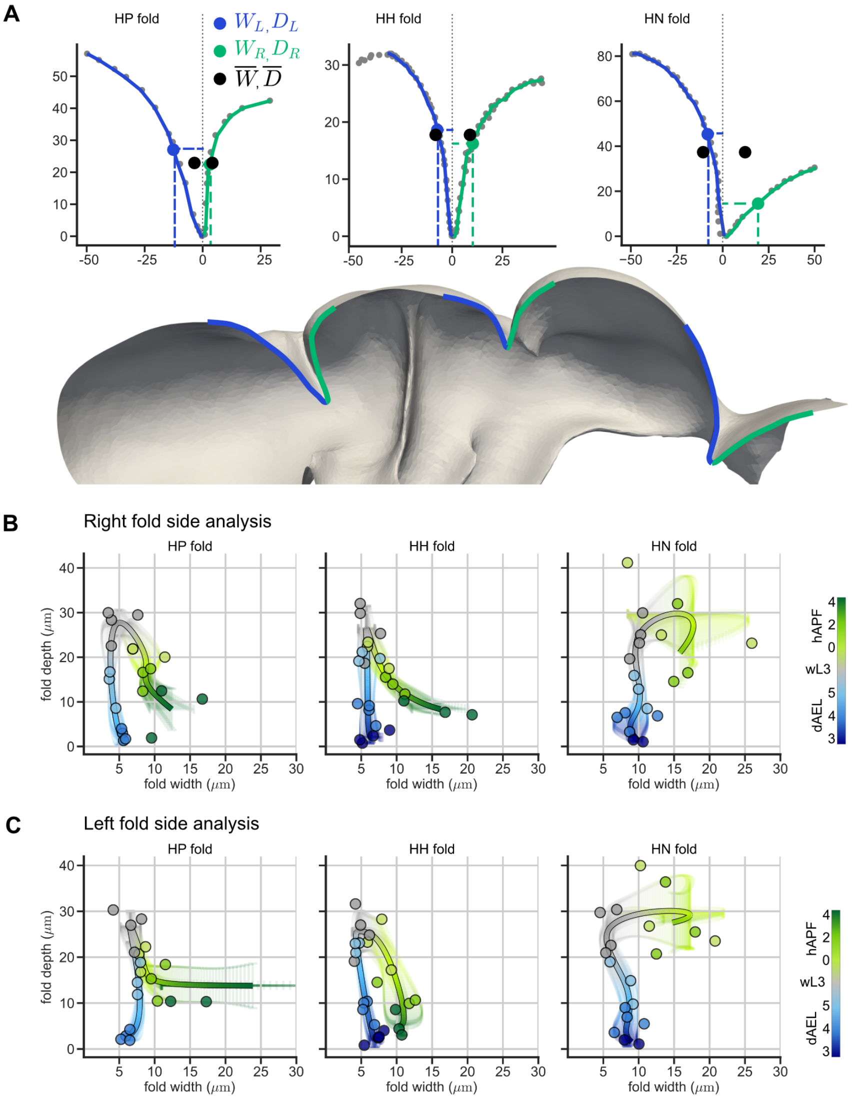


Figure S2: **A**, Plots of exemplary HP, HH, and HN fold cross-sections for a 2 hAPF WT wing disc, where grey points correspond to the mesh vertices and the solid lines show the left and right envelope. Dashed lines identify width ($W_L$ and $W_R$) and depth ($D_L$ and $D_R$) for each envelope, and the corresponding position on the envelope is highlighted with a circle. We then compute the cross-section width and depth ($\overline{W}$ and $\overline{D}$), shown by black circles, as a weighted average of the one-sided quantities, see Methods. Below: a cross-section through a 2 hAPF surface mesh, where the folds are indicated and colors indicate the direction (blue = left/ distal, green = right/ proximal). **B,C**, Plots of one-sided quantities per fold, with color indicating the developmental time. Data points indicate measured quantities of individual replicates, and the line shows the mean cubic spline interpolation over all data points. Error bars show the standard deviation of the mean spline. **B**, Width ($W_R$) and depth ($D_R$) of the right (proximal) side of the fold. **B**, Width ($W_L$) and depth ($D_L$) of the left (distal) side of the fold.

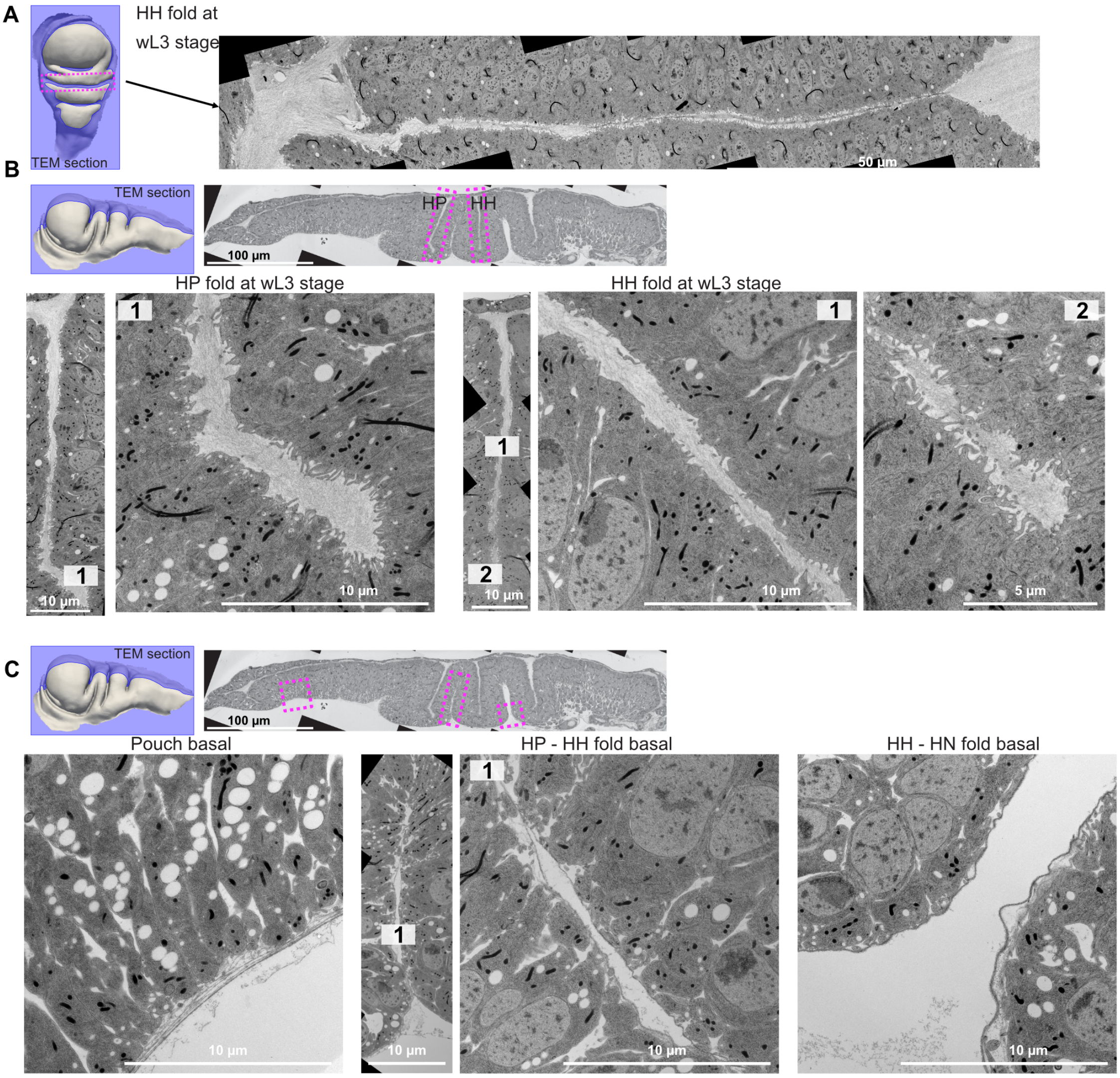


Figure S3: **A-C**, TEM of a wL3 wing disc. **A**, Top: Indication of the section orientation (wing disc surface in grey; TEM section in blue). The dashed box (magenta) highlights the HH fold region for which the TEM data is shown on the right. **B**, TEM of a cross-section along the long axis. Top: overview of the section with the HP and HH fold indicated by magenta boxes. Bottom row: left: overview of the HP fold, with the position of the higher magnification image on the right indicated (1) to show the apical extracellular space at the fold bottom; right: HH fold, with the higher magnification images in the middle of the fold (1) and at the base of the fold (2). **C**, Top shows the basal regions in the images below, indicated by magenta boxes. Left: basal view of the wing disc pouch; Middle: Region between the HP and HH fold, the overview image indicates the region (1) of the higher magnification. Right: Region between the HH and HN folds near the base of the folds.

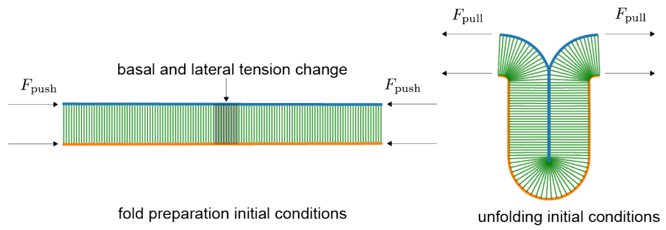


Figure S4: Left: initial tissue. To trigger fold formation, we change the basal and lateral tension and apply a boundary pushing force. Right: prepared fold after the equilibration stage. We apply various pulling forces and keep or remove adhesion to the mimic the presence or absence of aECM.

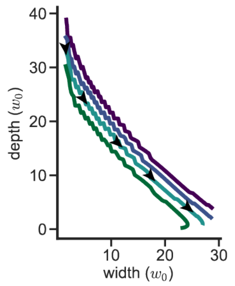


Figure S5: Depth vs width dynamics of simulations using various initial fold depths for a pulling force $F_{\mathrm{pull}} = 4F_0$ and no adhesion.

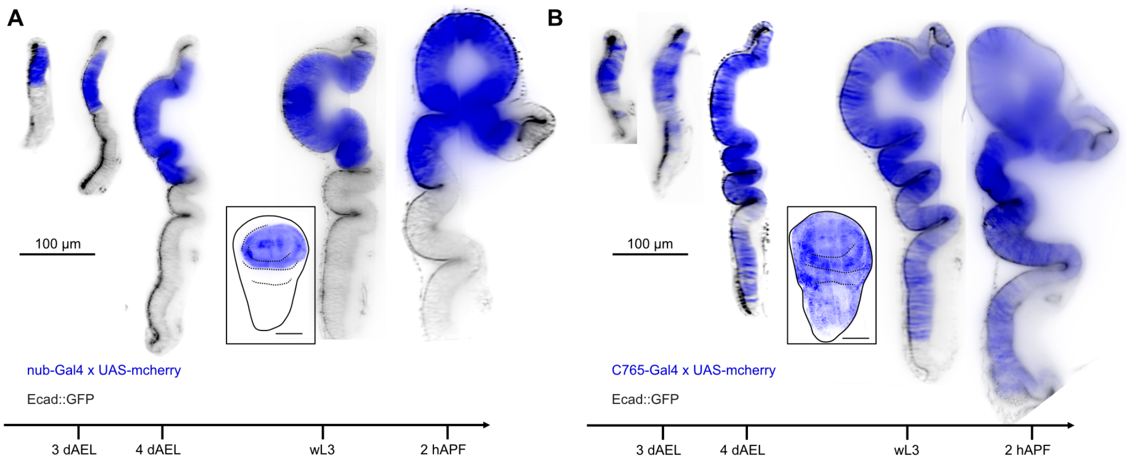


Figure S6: **A,B**, Long axis cross-section of different developmental stages showing the expression region of the Gal4 reporter (using UAS-mcherry) in blue and Ecad::GFP in black. The scale bar for all cross-sections is indicated on the left. The inset shows an apical view of a maximum intensity projection of UAS-mcherry at wL3 stage, with the wing disc outline (solid line) and the folds (dashed line) indicated (scale bar = 100 $\mu m$). **A**, nub-Gal4 driver; **B**, c765-Gal4 driver.

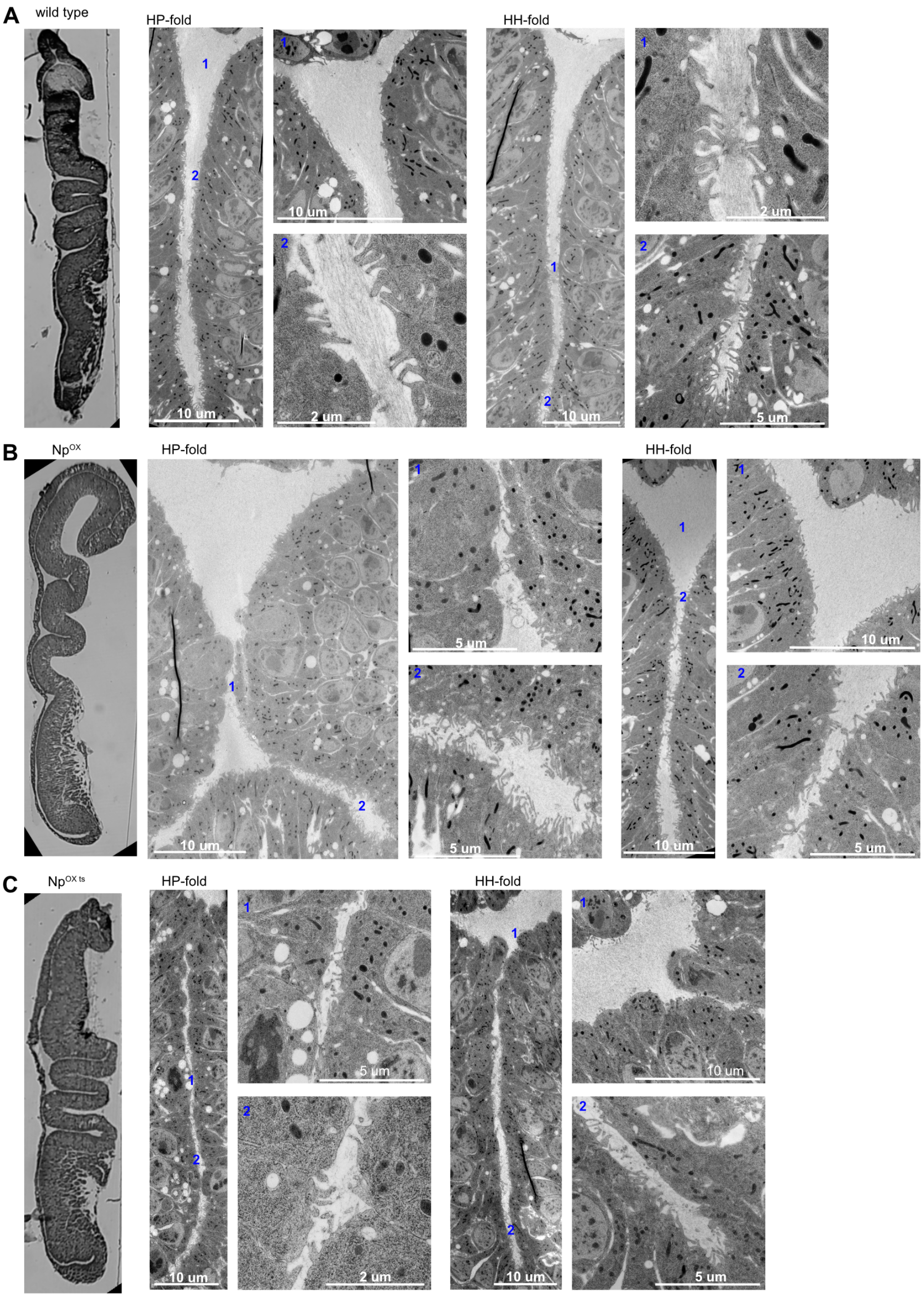


Figure S7: **A-C**, TEM images of long axis cross-sections at wL3 stage. Left: overview of each section, followed by an overview of the respective folds (HP and HH) and higher magnification images. The numbers indicate the positions of higher magnification images the within each fold. **A**, WT control, **B**, Np$^{OX}$, **C**, Np$^{OXts}$.

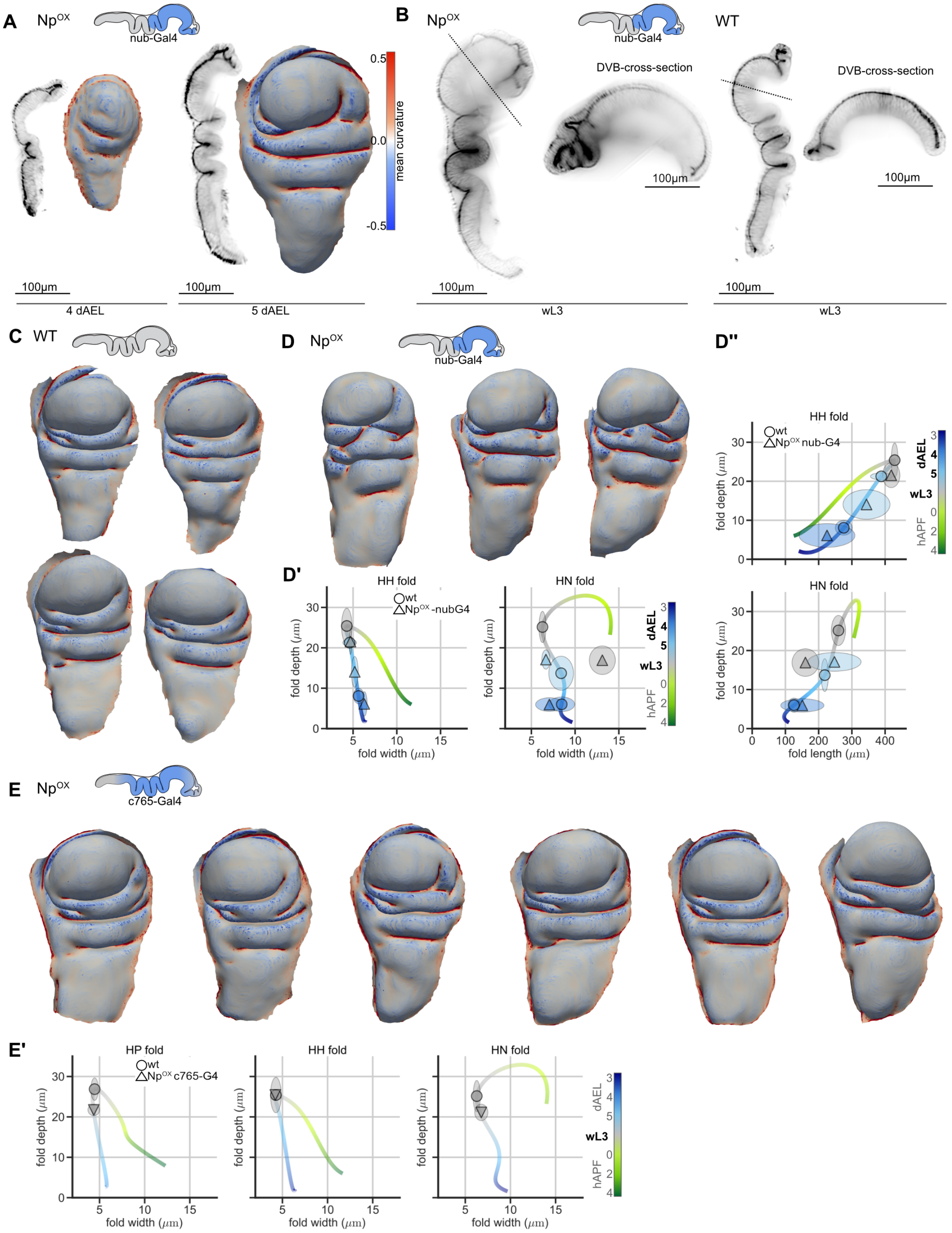


Figure S8: **A,C,D,E**, Apical surface meshes, anterior left, color coded by mean curvature ($\mu m^{-1}$); the color range indicated in **A** is the same for all meshes. **A-E**, Cartoons indicate the expression domain of the Gal4: A, B and D: nub-Gal4, C: WT, E: c765-Gal4. **A**, Long-axis cross-sections of wing discs labeled with Ecad::GFP and apical surface meshes of Np$^{OX}$ for 4 dAEL and 5 dAEL wing disc **B**, Long-axis and dorsal-ventral boundary (DVB) cross-sections for a wL3 Np$^{OX}$ and WT wing disc (label = Ecad::GFP). **C**, WT meshes at wL3 stage; **D**, Np$^{OX}$ meshes at wL3 stage; **D', D", E'**, Quantification of fold shape, plotted with the WT trajectory. Symbols: mean for WT and Np$^{OX}$; ellipse: extension along each axis corresponds to the 95% confidence interval of the mean. Color = developmental time. **D'**, Fold depth over width, **D"**, fold depth over length, **E'**, fold depth over width. **E, E'**, Meshes and quantification for the c765-Gal4 driver.

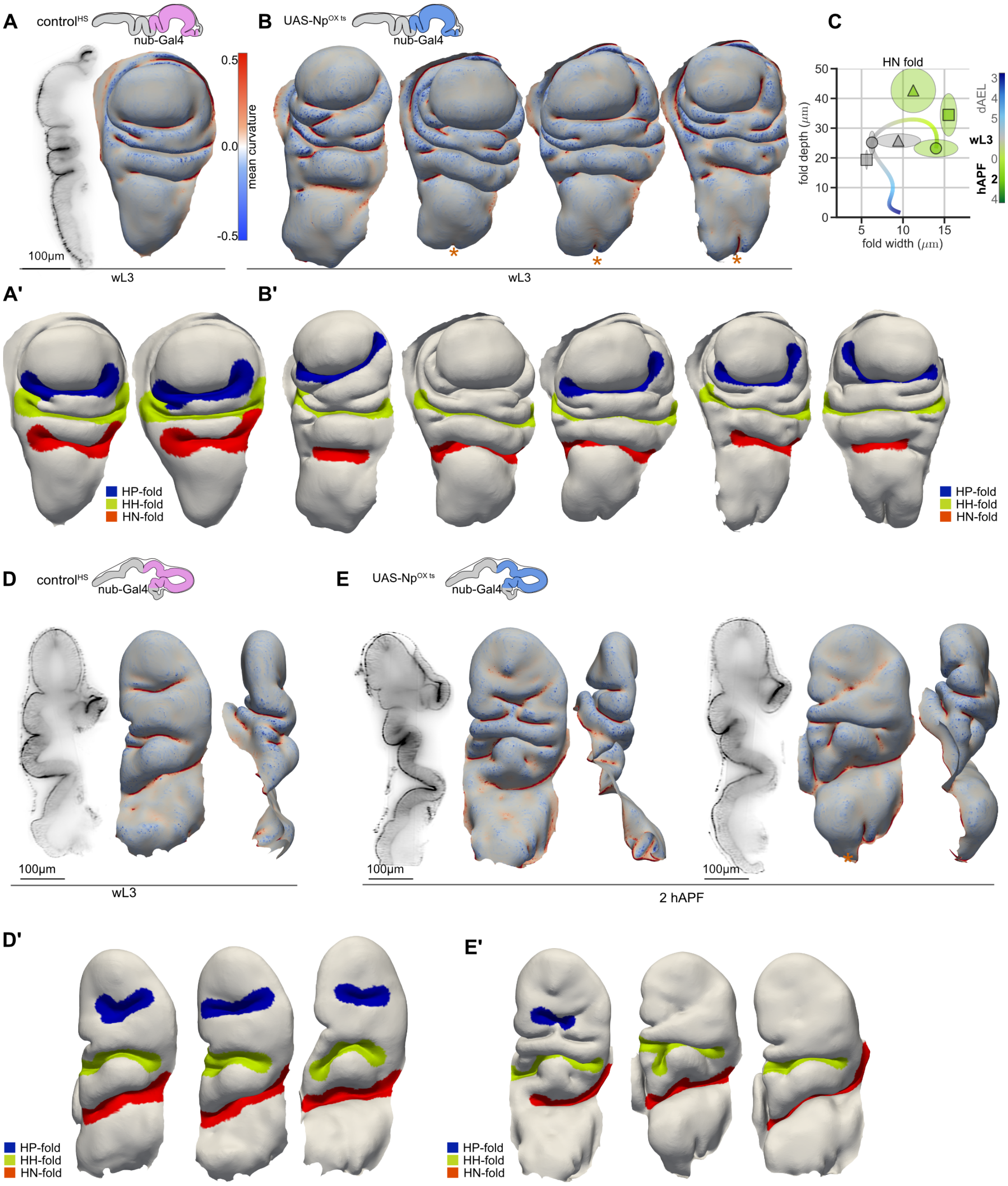


Figure S9: **A,B,D,E**, Apical surface meshes, head-on view: anterior left, color coded by mean curvature ($\mu m^{-1}$); the color range is the same for all meshes. Above: cartoons indicate the expression domain of the nub-Gal4, either combined with UAS-mcherry as HS control (control$^{HS}$, pink) or with Np$^{OXts}$ (blue). **A',B',D',E'**, Apical surface meshes with identified fold regions of the wing discs analyzed in C and Figure 4E, and shown in A,B,D,E and Figure 4D, head-on view: anterior left, identified fold regions are indicated in color. **A,D,E**, Long-axis cross-sections of wing discs labeled with Ecad::GFP next to the apical surface meshes. **A,A'**, wL3 stage of control$^{HS}$; **B,B'**, Meshes for Np$^{OXts}$; note that in one wing disc, the HP fold could not be identified using our method; **C**, Quantification of HN shape for Np$^{OXts}$, WT, and control$^{HS}$ (symbols: mean, ellipse: 95% confidence interval, color = developmental time). **D, E**, 2 hAPF stages with cross-section and meshes (left: head-on, right: anterior). **D,D'**, control$^{HS}$, **E,E'**, Np$^{OXts}$.

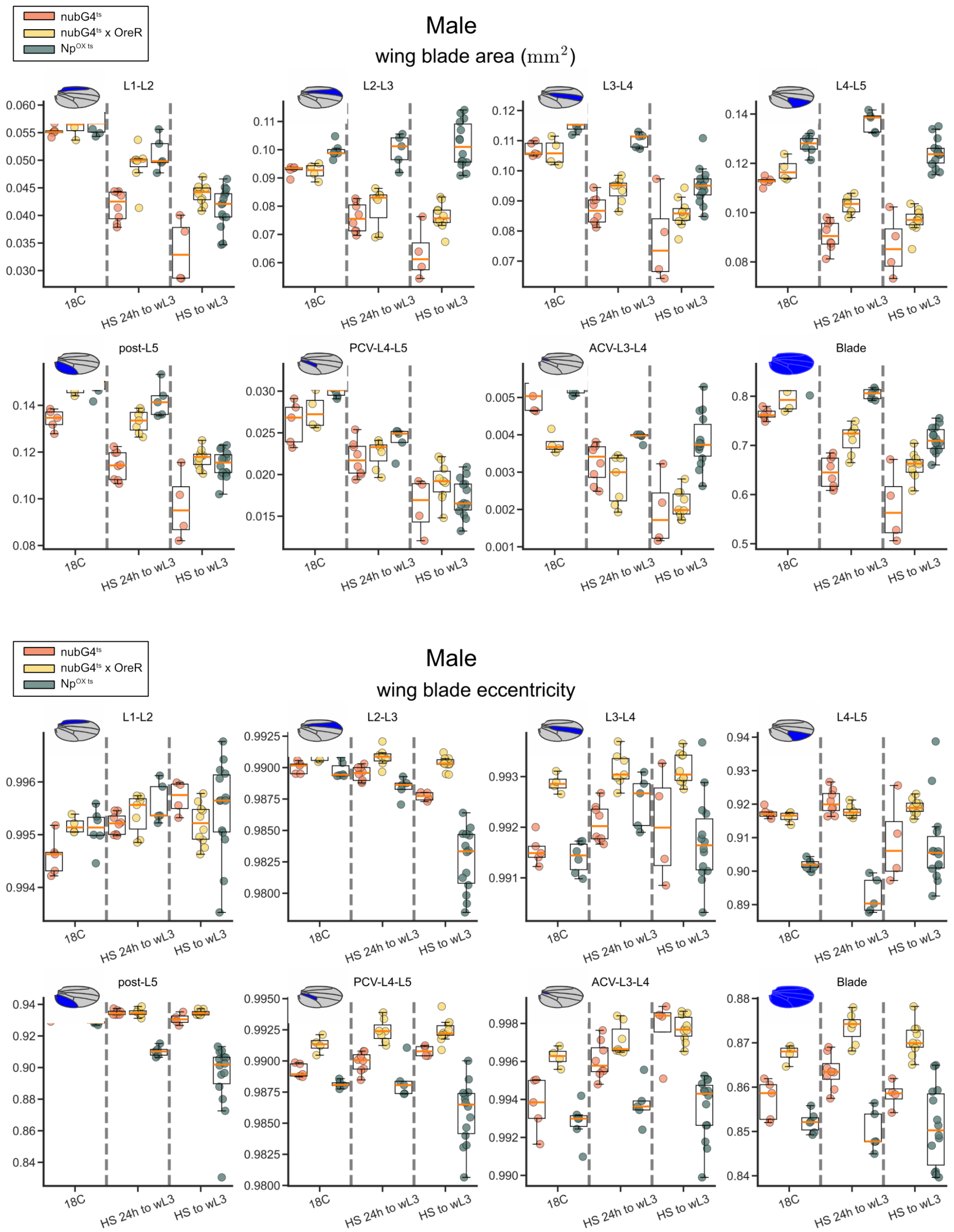


Figure S10: Analysis of adult wing shape for males of different genotypes and after temperature shifts. Conditions: No HS: raised at the permissive temperature for the inhibitor Gal80ts and therefore restricted expression of the Gal4 (18°C); 24h to wL3: raised at 18°C and then transiently shifted to 30°C for 24hrs prior to the wL3 stage; to wL3: shifted from egg laying to 30°C and grown there until wL3, when they were shifted and grown thereafter at 18°C. Genotypes: Control1: nub-Gal4$ts$, control2: nub-Gal4$ts$ x OreR, and Np$^{oxts}$. The analyzed regions are shown in the cartoons and indicated in the subtitles. Upper plots: area measurements in $mm^2$; bottom plots: eccentricity measurements. PCV = posterior crossvein, ACV = anterior crossvein.

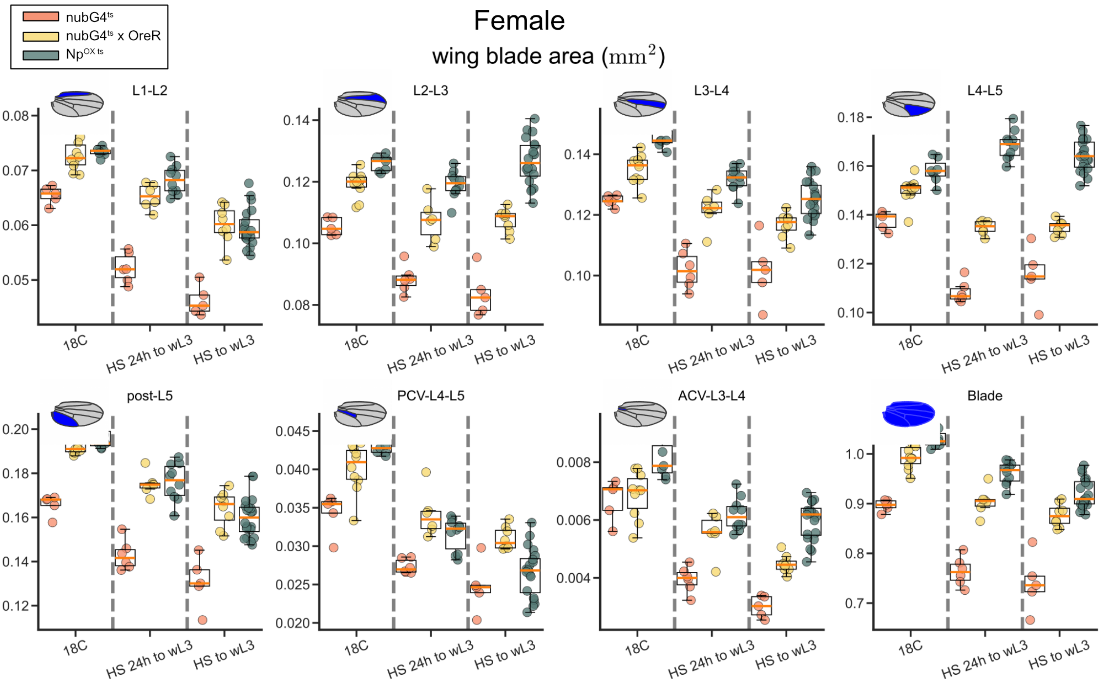


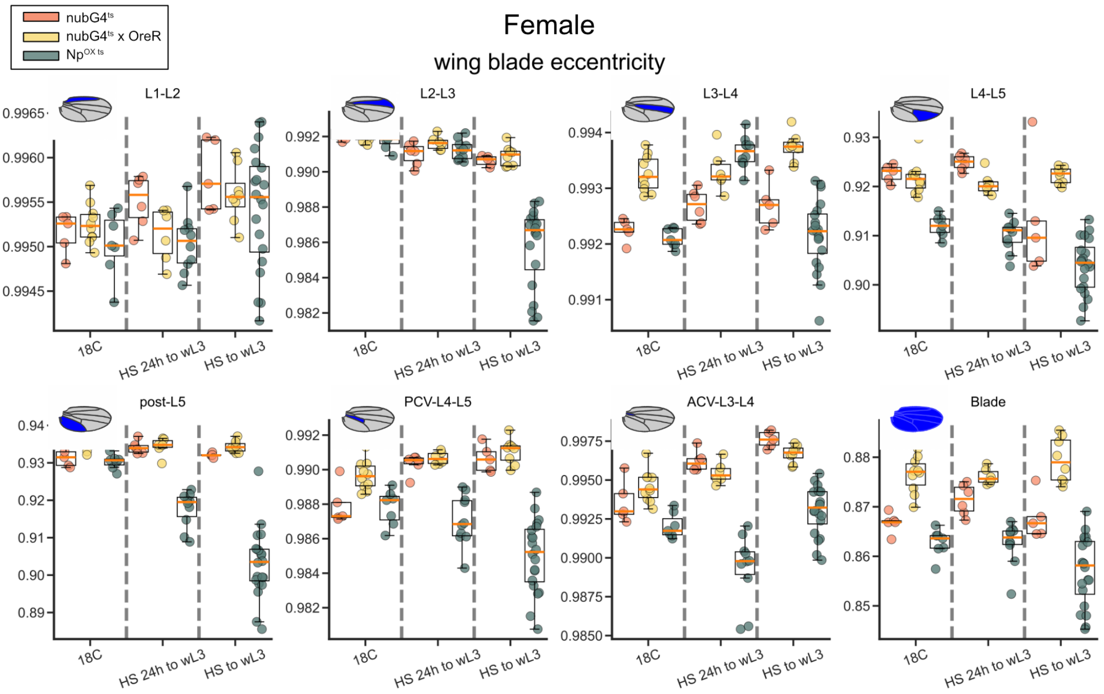


Figure S11: Analysis of adult wing shape for females of different genotypes and after temperature shifts. Conditions: No HS: raised at the permissive temperature for the inhibitor Gal80ts and therefore restricted expression of the Gal4 (18°C); 24h to wL3: raised at 18°C and then transiently shifted to 30°C for 24hrs prior to the wL3 stage; to wL3: shifted from egg laying to 30°C and grown there until wL3, when they were shifted and grown thereafter at 18°C. Genotypes: Control1: nub-Gal4$ts$, control2: nub-Gal4$ts$ x OreR, and Np$^{oxts}$. The analyzed regions are shown in the cartoons and indicated in the subtitles. Upper plots: area measurements in $mm^2$; bottom plots: eccentricity measurements. PCV = posterior crossvein, ACV = anterior crossvein.

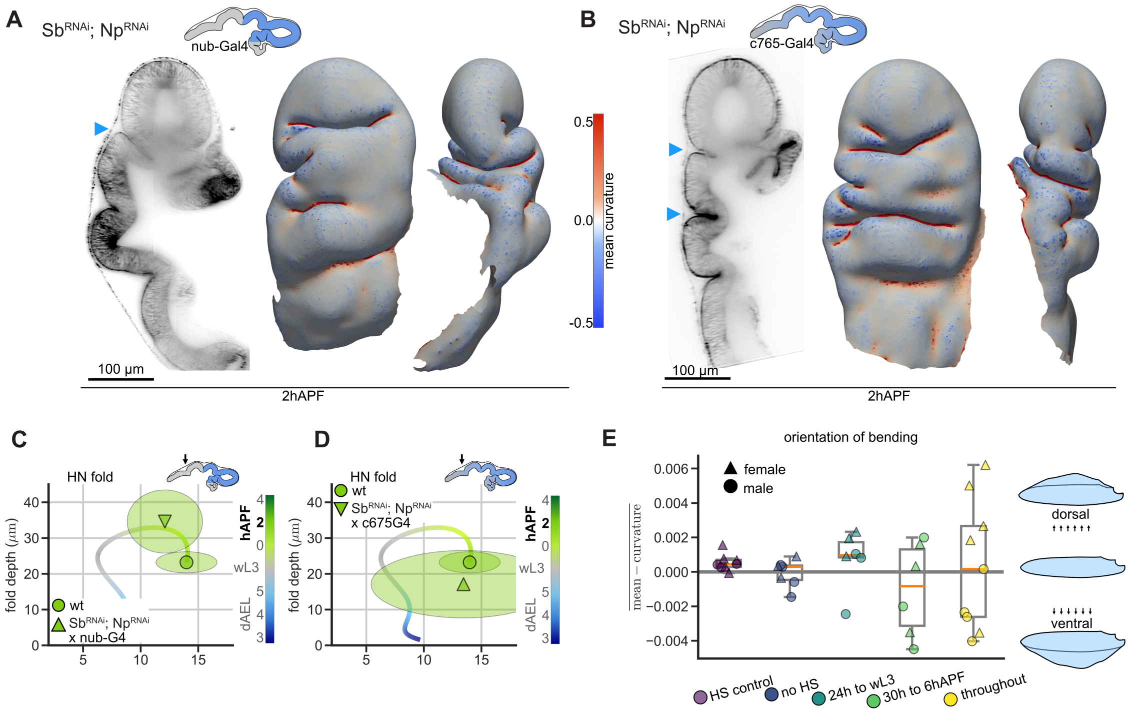


Figure S12: **A,B**, 2 hAPF wing discs of Sb$^{RNAi}$; Np$^{RNAi}$. Long-axis cross-sections of wing discs next to the apical surface meshes. Apical surface meshes, head-on view: anterior left, side view: anterior; color coded by mean curvature ($\mu m^{-1}$); the color range is the same for all meshes. Above: cartoons indicate the expression domain of nub-Gal4 and c765-Gal4 (blue) on 2hAPF wing discs. **A**, Sb$^{RNAi}$; Np$^{RNAi}$ with the nub-Gal4 driver. The cross-section image is Ecad::mTomato. **B**, Sb$^{RNAi}$; Np$^{RNAi}$ with the c765-Gal4 driver. The cross-section image is Ecad::GFP. **C,D**, Fold depth over width for the HN fold at 2hAPF (symbols: mean, ellipse: 95% confidence interval, color = developmental time). **C**, Sb$^{RNAi}$; Np$^{RNAi}$ with the nub-Gal4 driver. **D**, Sb$^{RNAi}$; Np$^{RNAi}$ with the c765-Gal4 driver. **E**, Average mean curvature ($\mu m^{-1}$) on adult wing meshes, taking only one side of the blade surface into account. A positive mean curvature indicates that the blade surface bends on average more toward the dorsal side; negative mean curvature indicates a bend in the ventral direction (see cartoon on the right). Analyzed are Sb$^{RNAi}$; Np$^{RNAi}$ with the nub-Gal4 driver in different heat shock (HS) conditions: HS control: control genotype with HS 24hrs prior to the wL3 stage, no HS: Sb$^{RNAi}$; Np$^{RNAi}$ nub-Gal4$^{ts}$, no HS, 24 h to wL3: Sb$^{RNAi}$; Np$^{RNAi}$ nub-Gal4$^{ts}$ HS for 24 h to wL3 stage, 30 h to 6 hAPF: Sb$^{RNAi}$; Np$^{RNAi}$ nub-Gal4$^{ts}$ HS for 30 h to 6 hAPF stage, throughout: Sb$^{RNAi}$; Np$^{RNAi}$ nub-Gal4.
